# HLA alleles associated with risk of ankylosing spondylitis and rheumatoid arthritis influence the gut microbiome

**DOI:** 10.1101/517813

**Authors:** Mark Asquith, Peter R. Sternes, Mary-Ellen Costello, Lisa Karstens, Sarah Diamond, Tammy M. Martin, Timothy D. Spector, Kim-Anh le Cao, James T. Rosenbaum, Matthew A. Brown

**Author notes:** Corresponding Author: Professor Matthew A. Brown, Institute of Health and Biomedical Innovation, Queensland University of Technology, Translational Research Institute, Princess Alexandra Hospital, Woolloongabba, Brisbane, Australia. These two authors contributed equally to this manuscript (James T. Rosenbaum, Matthew A. Brown).

## Abstract

**Objectives:** HLA alleles affect susceptibility to more than 100 diseases, but the mechanisms to account for these genotype-disease associations are largely unknown. HLA-alleles strongly influence predisposition to ankylosing spondylitis (AS) and rheumatoid arthritis (RA). Both AS and RA patients have discrete intestinal and faecal microbiome signatures. Whether these changes are cause or consequence of the diseases themselves is unclear. To distinguish these possibilities, we examine the effect of *HLA-B27* and *HLA-DRB1* RA-risk alleles on the composition of the intestinal microbiome in healthy individuals.

**Methods:** 568 samples from 6 intestinal sites were collected from 107 otherwise healthy unrelated subjects and stool samples from 696 twin pairs from the TwinsUK cohort. Microbiome profiling was performed using sequencing of the 16S rRNA bacterial marker gene. All patients were genotyped using the Illumina CoreExome SNP microarray, and HLA genotypes were imputed from these data.

**Results:** Association was observed between *HLA-B27* genotype, and RA-risk *HLA-DRB1* alleles, and overall microbial composition (P=0.0002 and P=0.00001 respectively). These associations were replicated in the TwinsUK cohort stool samples (P=0.023 and P=0.033 respectively).

**Conclusions:** This study shows that the changes in intestinal microbiome composition seen in AS and RA are at least partially due to effects of *HLA-B27* and –*DRB1* on the gut microbiome. These findings support the hypothesis that HLA alleles operate to cause or increase the risk of these diseases through interaction with the intestinal microbiome, and suggest that therapies targeting the microbiome may be effective in their prevention and/or treatment.

## INTRODUCTION

HLA molecules affect susceptibility to many diseases, but in the majority of cases the mechanism by which HLA molecules predispose to disease remains a mystery. The risks of developing both ankylosing spondylitis (AS) and rheumatoid arthritis are primarily driven by genetic effects, with heritability >90% (1, 2) for AS, and 53-68% for RA (3, 4). In both diseases HLA alleles are the major susceptibility factors, with AS being strongly associated with *HLA-B27*, and RA with *HLA-DRB1* ‘shared-epitope’ (SE) alleles.

Particularly in AS, there is strong evidence of a role for gut disease in disease pathogenesis. Up to an estimated 70% of AS patients have either clinical or subclinical gut disease, suggesting that intestinal inflammation may play a role in disease pathogenesis (5, 6). Increased gut permeability has been demonstrated in both AS patients and their first-degree relatives compared with unrelated healthy controls (7-11). Crohn’s disease (CD) is closely related to AS with a similar prevalence and high heritability. The two commonly co-occur with an estimated ∼5% of AS patients developing CD, and 4-10% of CD patients developing AS (12, 13). Strong co-familiality (14), and the extensive sharing of genetic factors between AS and inflammatory bowel disease (IBD) (15, 16) suggests that they have a shared aetiopathogenesis. This is consistent with the hypothesis that gut derived immune cells or microbial products may contribute to spondyloarthritic inflammation (17-19).

Using 16S rRNA community profiling we have previously demonstrated that AS cases have a discrete intestinal microbial signature in the terminal ileum (TI) compared with healthy controls (HC) (P<0.001) (20), a finding that has subsequently been confirmed by other studies (21, 22). We have also demonstrated that dysbiosis is an early feature of disease in *HLA-B27* transgenic rats, preceding the onset of clinical disease in the gut or joints (23). Similarly, RA cases have also been shown to have gut dysbiosis (24, 25), and animal models of RA such as collagen-induced arthritis have been shown to be influenced by the gut microbiome (26, 27). In these studies it is difficult to distinguish between effects of the immunological processes going on in the intestinal wall in cases, and the effects of treatments on the intestinal microbiome, from potential effects of the gut microbiome on the disease.

The role of the host genetics in shaping intestinal microbial community composition in humans is unclear. In animal models, host gene deletions have been shown to result in shifts in microbiota composition (28). In addition, a recent quantitative trait locus mapping study in an inter-cross murine model, linked specific genetic polymorphisms with microbial abundances (29). Large scale studies in twins (n=1126 twin pairs) have demonstrated that of 945 widely shared taxa, 8.8% showed significant heritability, with some taxa having heritability of >40% (e.g. family *Christensenellaceae*, heritability 42%) (30).

Further studies are needed into whether the changes in intestinal microbial composition are due to host genetics, and how this affects the overall function of the gut microbiome in cases, including how the microbiome then goes on to shape the immune response and influence inflammation. In AS, given the strong association of *HLA-B27*, the hypothesis has been raised that *HLA-B27* induces AS by effects on the gut microbiome, in turn driving spondyloarthritis and inducing immunological processes such as IL-23 production (31, 32). Further experiments comparing the intestinal microbiome of *HLA-B27* negative and positive patients would shed light of the influence of *HLA-B27* on overall intestinal microbiome composition, particularly given the work in *HLA-B27* transgenic rats showing that *HLA- B27* was associated with altered ileal, caecal, colonic and fecal microbiota (23, 33, 34). Similar theories have been proposed with regard to interaction between the gut microbiome and the immunological processes that drive RA (reviewed in (35)).

In this study we investigated if AS and RA-associated HLA alleles influence the gut microbiome in healthy individuals, to support the hypothesis that they influence the risk of developing AS and RA through effects on the gut microbiome.

## METHODS

### Human subjects

A total of 107 subjects, aged 40-75, predominately Caucasian (∼90%), typically following an omnivorous diet (∼95%) and were undergoing routine colorectal cancer screening at Oregon Health & Science University’s Center for Health and Healing were included in this study. Individuals were excluded if they had a personal history of inflammatory bowel disease or colon cancer, prior bowel or intestinal surgery or were pregnant. All subjects underwent a standard polyethylene glycol bowel prep the day prior to their colonoscopy procedure. During the procedure, biopsies were collected for research purposes from the terminal ileum or other tissue sites as indicated. Subjects were instructed to collect a stool sample on a sterile swab at home, just prior to starting their bowel prep procedure. Stool samples were brought to the colonoscopy appointment at room temperature. All samples (biopsies and fecal swabs) were placed at 4°C in the clinic and transported to the lab within 2 hours of the colonoscopy procedure, where they were snap frozen and stored at −80°C prior to processing. Patient samples were obtained over a 24-month period.

Ethical approval for this study was obtained from the Oregon Health & Science University Institutional Review Board. Written informed consent was obtained from all subjects. This study was performed subject to all applicable U.S. Federal and State regulations.

### TwinsUK

All work involving human subjects was approved by the Cornell University IRB (Protocol ID 1108002388). Matched genotyped and stool samples were collected from 1392 twins. Genotyping, 16S rRNA amplicon sequencing, filtering and analysis were performed as described in Goodrich *et al.,* 2014 (36).

### 16S rRNA amplicon sequencing and analysis

568 stool and biopsy samples across 107 individuals were extracted and amplified for the bacterial marker gene 16S rRNA as previously described (20). Samples were demultiplexed and filtered for quality using the online platform BaseSpace (http://basespace.illumina.com). Paired end reads were joined, quality filtered and analysed using Quantitative Insights Into Microbial Ecology (QIIME) v1.9.1 (37). Operational taxonomy units (OTU) were picked against a closed reference and taxonomy was assigned using the Greengenes database (gg_13_8) (38), clustered at 97% similarity by UCLUST (39) and low abundance OTUs were removed (<0.01%).

### Data visualization and statistical analysis

Multidimensional data visualisation was conducted using a sparse partial least squares discriminant analysis (sPLSDA) on centered log ratio transformed data, as implemented in R as part of the MixOmics package v6.3.1 (40). Association of the microbial composition with metadata of interest was conducted using a PERMANOVA test as part of vegan v2.4-5 (41) on arcsine square root transformed data at species level, taking into account individual identity where multiple sites per individual were co-analysed, as well as the sources of covariation such as BMI and gender. Alpha diversity was calculated at species level using the rarefy function as implemented in vegan v2.4-5 and differences were evaluated using a Wilcoxon rank-sum test. The metagenome functional content was predicted using PICRUSt v1.1.3 (42) and the resulting predictions were mapped to KEGG pathways using HUMAnN2 v0.11.1 (43) Differential abundance of bacterial taxa and KEGG pathways were tested for significance using MaAsLin v0.0.5 (44).

### Genotyping

DNA was extracted from mucosal biopsies and stool samples, and genotyped using Illumina CoreExome SNP microarrays according to standard protocols. Bead intensity data were processed and normalized for each sample, and genotypes called using Genome Studio (Illumina). We imputed *HLA-B* genotypes using SNP2HLA **(45)**, as previously reported **(46)**. The distribution of *HLA-B27* and *HLA-DRB1* RA-risk, -protective and –neutral subtypes is available in Supplementary Table 1.

## RESULTS

16S rRNA profiling and SNP array genotyping was successfully completed for 107 individuals (61 female, 46 male) involving a total of 564 biopsy samples (see Table 1).

**Table 1:** Number of samples and *HLA-B27* and *HLA-DRB1* shared epitope allele status by site. Note that different subjects had different numbers of samples obtained, and at no individual site did all subjects have samples obtained.

| Site | Total | Female | Male | HLA-B27<br>Negative | HLA-B27<br>Positive | HLA-DRB1<br>Risk<br>Genotype | HLA-DRB1<br>Protective<br>Genotype | HLA-DRB1<br>Neutral<br>Genotype |
| --- | --- | --- | --- | --- | --- | --- | --- | --- |
| Cecum | 103 | 59 | 44 | 93 | 10 | 34 | 8 | 47 |
| Ileum | 90 | 51 | 39 | 80 | 10 | 36 | 8 | 45 |
| Left Colon | 100 | 57 | 43 | 90 | 10 | 33 | 7 | 47 |
| Rectum | 91 | 53 | 38 | 81 | 10 | 33 | 7 | 41 |
| Right Colon | 97 | 57 | 40 | 87 | 10 | 33 | 8 | 45 |
| Stool | 83 | 46 | 37 | 73 | 10 | 29 | 8 | 36 |

We studied the effect of BMI, gender and sampling site on the gut microbiome to identify relevant covariates for analysis of AS-associated genes and their association with the gut microbiome. Considering sample site, striking differences were observed, particularly between the stool samples and mucosal samples (Figure 1A, P<0.0001). Excluding stool samples, marked difference was still observed between sites (P<0.0001), but it can be observed that this is mainly driven by differences of the ileal samples from the colonic mucosal samples (left and right colon, cecum, rectum), which largely clustered together (Figure 1B).

**Figure 1:**
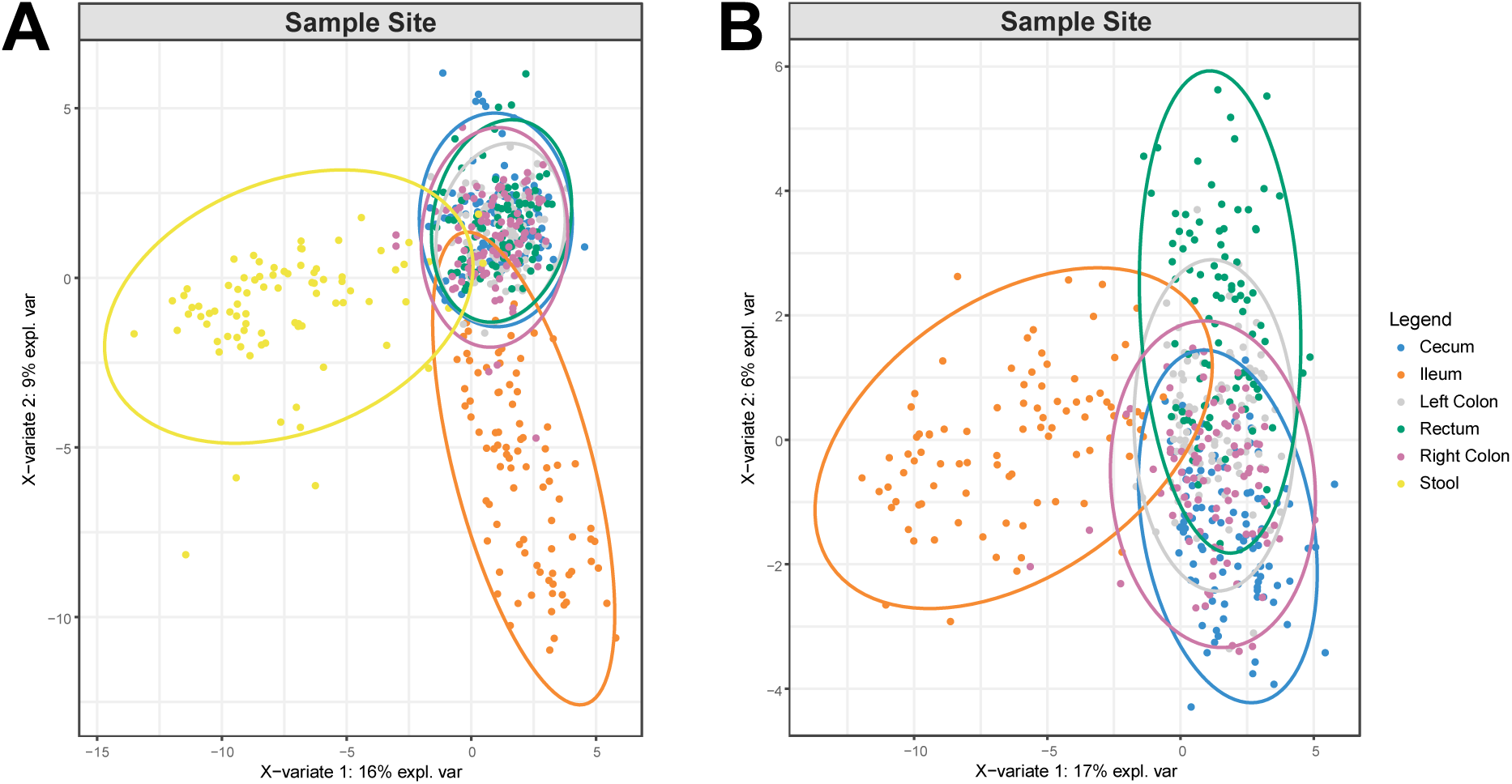
sPLSDA comparing the microbiome composition at various sample sites, showing **A.** marked difference of stool/luminal site compared with all other sites, which are mucosal, and **B.** in the absence of stool samples, the ileal site remains distinct from colonic sites. A PCA plot of these results is available in Supplementary Figure 1.

Stool samples are much more convenient to obtain than ileal or colonic mucosal samples, which require an endoscopic procedure for collection. Given the prior evidence of primarily ileal inflammation in AS (5), we were interested in the relationship between the ileal and stool microbiome. In this comparison marked differences were observed between sites, though with some overlap seen on the sPLSDA plot (Supplementary Figure 2, P<0.0001).

Several studies have noted an increase (47), decrease (20, 21) or no change (48) in alpha diversity metrics for AS cases, and an increase (22) or decrease (49) in alpha diversity for RA cases. In the current study, calculation of rarefied species richness revealed that carriage of *HLA-B27* and *HLA-DRB1* alleles was not associated with differences in alpha diversity, except for stool samples for which carriage of *HLA-DRB1* RA-risk alleles was associated an increased alpha diversity across both cohorts (Figure 2).

**Figure 2:**
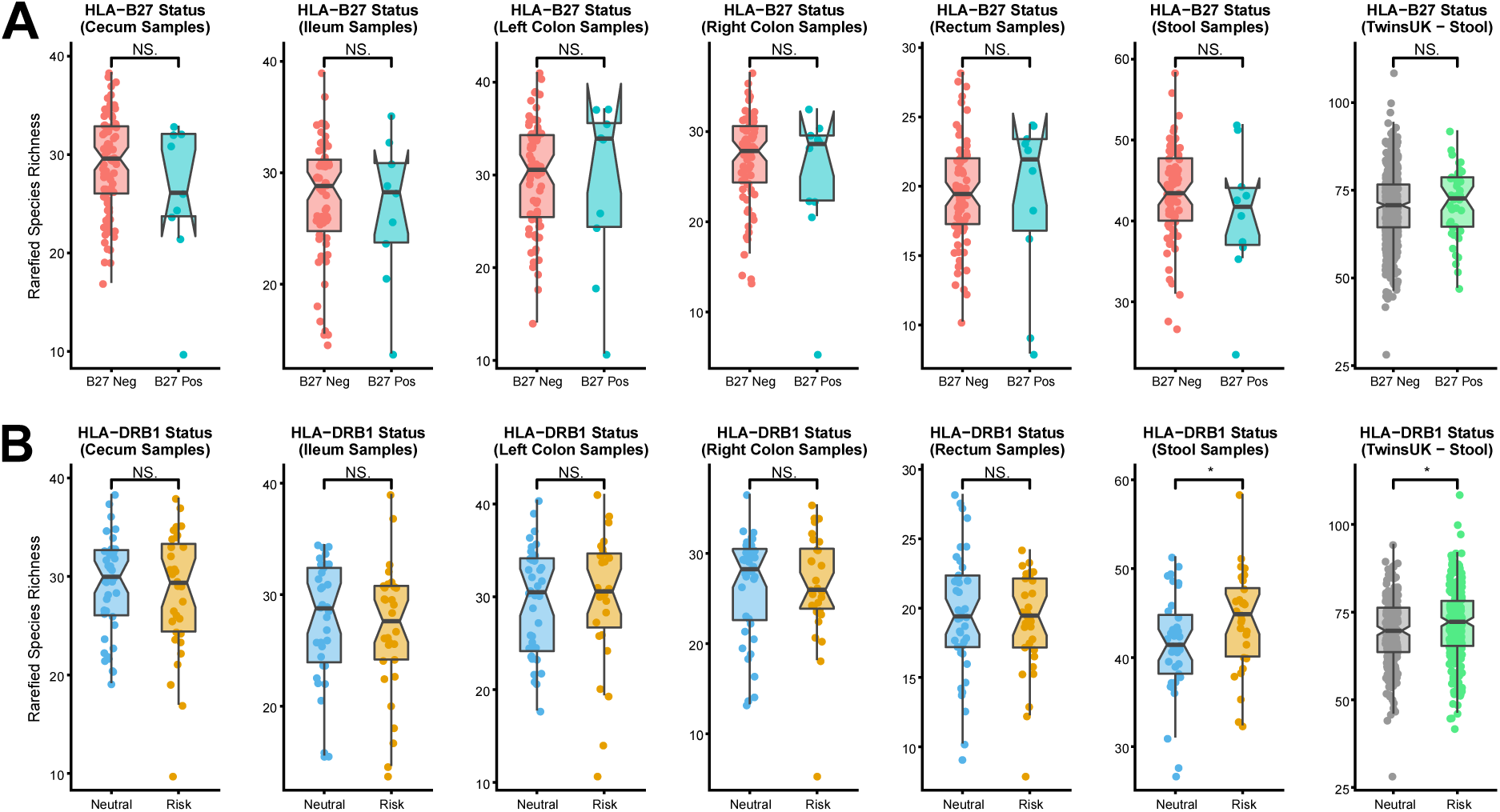
Alpha diversity across each sampling site, and in the TwinsUK cohort **A.** Alpha diversity according *HLA-B27* status. **B.** Alpha diversity according to *HLA-DRB1* status, revealing increased alpha diversity in stool samples of both cohorts.

Considering beta diversity via sPLSDA and PERMANOVA, significant association of BMI category was seen with microbiome composition (P=0.0022)(Supplementary Figure 3A). This appears to be driven particularly by the difference of underweight individuals (BMI<18.5) compared with other BMI categories. Removing underweight samples from the analysis, a non-significant trend of association of BMI category with microbiome composition is seen (P=0.078)(Supplementary Figure 3B), consistent with previous reports (50-52).

Given the marked gender biases in RA and AS, and evidence in mice that gender related hormonal differences are associated with differences in the intestinal microbiome (53, 54), we sought to evaluate the influence of gender on the microbiome in this cohort. Whilst substantial overlap between males and females was evident (Supplementary Figure 4), significant difference between genders in microbiome composition was observed (considering all sites, P=0.0004). Considering indicator species, a significant reduction in carriage of *Prevotella* genus in males was observed (P=0.005).

Controlling for BMI and gender, significant differentiation of the microbiome was identified in individuals carrying *HLA-B27* or RA-risk *HLA-DRB1* alleles (PERMANOVA P=0.002 and P=0.0001, respectively)(Figures 3A and 3B). Despite significant differentiation in terms of beta diversity, there was typically no difference in alpha diversity (Figure 2), indicating that the underlying host genetics may affect the overall composition of the microbiome, but not the overall species diversity. In the TwinsUK cohort, consisting of stool samples, and studying one twin drawn randomly from each twin pair, association with *HLA-B27* and RA-risk *HLA-DRB1* alleles was also observed (P=0.023 and P=0.033 respectively, Figure 3C). Study of the alternate twin from each pair revealed consistent findings. Whether the observed differences in taxonomic and functional composition are consistent between the two cohorts remains an open-ended question as they are confounded by differences in the experimental approach and the surveyed population.

**Figure 3:**
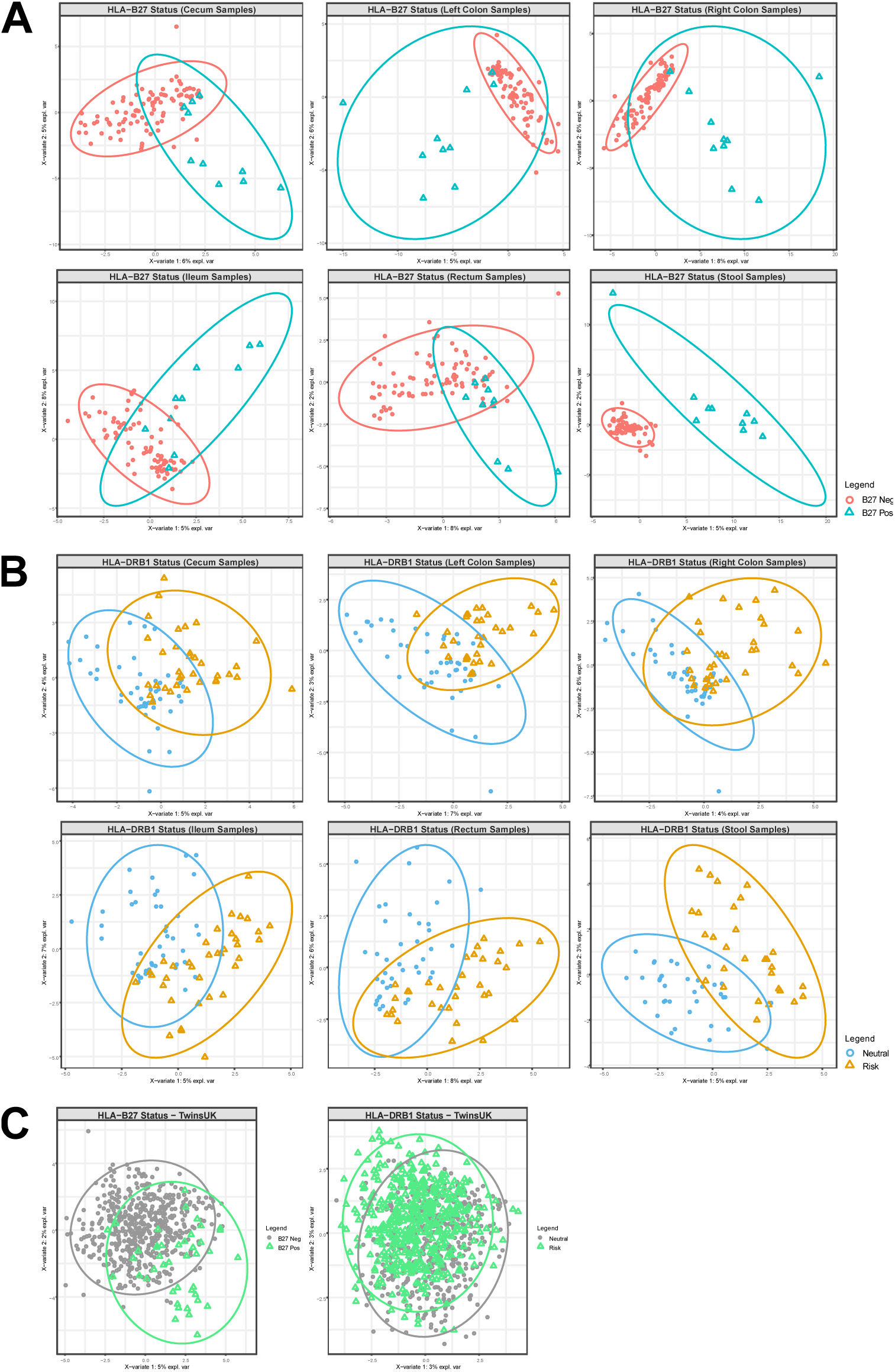
A. sPLSDA comparing the microbiome composition of *HLA-B27* positive and negative individuals across each sampling site. Considering all sampling sites and accounting for repeated sampling, significant differentiation of the microbiome was observed (PERMANOVA P=0.002). **B.** sPLSDA comparing individuals carrying the *HLA-DRB1* RA-risk and -neutral genotypes across each sampling site. Considering all sites and accounting for repeated sampling, significant differentiation of the microbiome was observed (PERMANOVA P=0.0001). **C.** sPLSDA plot comparing *HLA-B27* positive and negative twins (one twin randomly selected from each twin pair, PERMANOVA P=0.023), and *HLA-DRB1* risk and neutral genotypes (one twin randomly selected from each twin pair, PERMANOVA P=0.033). PCA plots of these results are available in Supplementary Figure 5.

We tested whether HLA-B alleles associated with AS were also associated with gut microbial profiles. The association of HLA-B alleles with AS is complex, with risk associations observed with *HLA-B27, -B13, -B40, -B47* and *–B51*, and protective associations with *HLA-B7* and *–B57* (55). Of these, only *HLA-B27* showed statistically significant association with microbiome profile across both cohorts. Differences in the microbiome composition were more pronounced when comparing risk-associated alleles to protective alleles. For example, when focusing on a subset of data (ileal samples), marginal differentiation for *–B27* (P=0.16) and no differentiation for *–B7* (P=0.61) was observed, potentially highlighting sample size constraints. However, direct comparison of *–B27* to *–B7* revealed significant differentiation (P=0.008).

*HLA-B27*-positive subjects exhibited reduced carriage (P<0.05) of *Bacterioides ovatus* across multiple sites (ileum, cecum, left colon, right colon and stool), as well as *Blautia obeum* (left colon and right colon) and *Dorea formicigenerans* (rectum and stool). Increased carriage of a *Roseburia* species was observed across multiple sites (left colon, right colon, rectum and stool) and family *Neisseriaceae* (cecum and ileum). For subjects with RA-risk *HLA-DRB1* alleles, numerous taxonomic groups were enriched across multiple sites, notably a *Lachnospiraceae* species (ileum, cecum, left colon, right colon and rectum), a *Clostridiaceae* species (left colon, right colon, rectum and stool) *Bifidobacterium longum* (cecum, right colon and rectum), amongst many others. Enrichment of *Ruminococcus gnavus* was also observed in the ileum of subjects carrying risk alleles. A full list of differently abundant taxa according to *HLA-B27* and *HLA-DRB1* status are available in Supplementary Tables 2 and 3, respectively. Interestingly, when accounting for false discovery rate, no single taxa was significantly associated with the investigated genotypes, indicating that community-level differences detectable via PERMANOVA may be driven by subtle changes in a high number of taxa, as opposed to marked changes in a select few.

Considering the inferred metabolic profiles for *HLA-B27* positive and negative subjects, we observed significant differences (P<0.05) across multiple sites for numerous KEGG pathways (Supplementary Table 4). Examples include flagellar assembly (ileum, cecum, left colon, right colon and rectum), alanine metabolism (cecum, ileum, left colon, and right colon), lysine biosynthesis (left and right colon) and degradation (ileum, rectum and stool) and secondary bile acid biosynthesis (ileum and stool). For the RA-risk alleles (*HLA-DRB1*), numerous differences in KEGG pathways were observed (Supplementary Table 5). Examples include thiamine metabolism, the citric acid cycle, lipopolysaccharide biosynthesis, glycerolipid metabolism biosynthesis of ansamycins, RNA transport and bacterial chemotaxis, all of which were differentially abundant across every tissue site biopsied.

## DISCUSSION

In this study we have demonstrated for the first time that in the absence of disease or treatment, *HLA-B27* and *HLA-DRB1* have significant effects on the gut microbiome in humans. This is consistent with *HLA-DRB1-*associated observations in mice (56) and the effect of *HLA-DRB1* alleles upon *Prevotella copri* abundance in humans (24). This extends previous demonstrations that AS and RA are characterized by intestinal dysbiosis by confirming that this is at least in part due to the effects of the major genetic risk factors for AS and RA, *HLA-B27* and *HLA-DRB1 risk* alleles, respectively.

We demonstrate a clear distinction in microbiome profile between luminal stool samples and mucosal samples, as well as between mucosal samples from different intestinal sites. Of particular note, marked difference was observed between ileal and stool samples. These findings contrast a previous smaller study, which may not have observed a difference between ileal and colonic biopsies due to sample size considerations (48). Many studies of the influence of gut microbiome focus on stool samples, as they are easier to obtain than mucosal samples. The existence of gut inflammation, particularly involving the ileum, in AS cases has been well documented. Therefore, our findings suggest that studies of the microbiome in AS and RA, particularly where the aim is to identify the key species or genetic elements driving or protecting from the disease, should use samples that reflect the site of inflammation (i.e. at least in AS, ideally the ileal microbiome). As the microbiome profile of stool samples do not closely correlate with the ileal microbiome, they would not appear to be an optimal sample to study, although studying IgA coated bacteria isolated from stool samples may prove more informative (57, 58).

Following our initial study, three further studies have now reported on the difference in gut microbial composition in AS cases and controls. Tito et al (48) in a study of 27 spondyloarthritis patients (i.e. not necessarily AS) and 15 healthy controls using 16S rRNA profiling report association of carriage of *Dialister* in ileal or colonic mucosal biopsies with disease activity assessed by the self-reported questionnaire the Bath Ankylosing Spondylitis Disease Activity Index (BASDAI), and Ankylosing Spondylitis Disease Activity Score (ASDAS). We did not observe *Dialister* in our study and therefore cannot comment on whether it is associated with *HLA-B27* carriage. Tito et al did not observe association of the gut microbiome with *HLA-B27* carriage, but the sample size, particularly in healthy controls, was too small to exclude other than a large effect. Wen et al used shotgun sequencing of stool samples from in 97 Chinese AS cases and 114 healthy controls, and reported significant dysbiosis in the AS cases (21). Breban et al (22) used 16S rRNA profiling of the stool microbiome to study 87 +-patients with axial spondyloarthritis (42 with AS), 69 healthy controls and 28 rheumatoid arthritis patients. They also report evidence of intestinal dysbiosis in the spondyloarthritis patients, and report correlation of *Ruminococcus gnavus* carriage with BASDAI. Whilst we did not observe an association with the carriage of *HLA-B27, Ruminococcus gnavus* was noted to be enriched in the ileum of individuals carrying the *HLA-DRB1* RA-risk alleles (Supplementary Table 3). In a comparison of *HLA-B27* positive and negative siblings (n=22 and 21 respectively), no difference in microbial composition was noted overall, but *HLA-B27* positive siblings had increased carriage of the *Microcaccaceae* family (including the species *Rothia mucilaginosa* within it), several *Blautia* and *Ruminococcus* species, and of *Egerthella lenta*. They also observed a reduced carriage of *Bifidobacterium* and *Odoribacter* species. Of these we also see reduction in *Blautia obeum*. Although we did not find dysbiotic changes that were shared with these specific taxa, we note the enrichment of genera within the Lachospiraceae-Ruminococcaceae grouping in *HLA-B27* carriers was a shared feature of our studies; *Roseburia* and *Ruminococcus* by Breban et al (22) and *Roseburia, Blautia, Dorea* and *Oscillospira* in our current study. These bacteria are known to be enriched within the intestinal mucosa (59), and are plausibly more immunogenic as a result (60). The differences observed between these studies may relate to analytical differences such as handling of covariates, disease definition, sample site studied, ethnicity and diet, and the different methods employed to profile the microbiome. Our study also confirms the significant effect of gender and BMI category on gut microbial profiles, suggesting that future studies should control for these covariates. Consistent with a recent study which examined the effect of the host’s genetics upon the microbiome of 1,046 healthy individuals (61), numerous correlations between specific bacterial taxa and the host’s genotype do not remain significant following correction for false discovery rate, thus indicating that HLA molecules may have a more generalized effect upon microbiome composition as opposed to a marked effect upon specific taxa. Despite this, we note that many of the P < 0.05 associations occurred across multiple tissue sites. Whilst the chance of a false positive at a single site might be relatively high, the chances of finding the same association across multiple sites decreases exponentially, indicating that the results are less likely to be spurious. Another possibility is that differences in microbial gene content, not necessarily specific taxa, may be more significant. In the current study, the microbiome’s predicted gene content was extrapolated from the underlying taxonomy, therefore utilization of whole genome sequencing metagenomics (a.k.a. shotgun metagenomics) to directly profile genetic composition may prove fruitful. This will be the focus of subsequent studies.

HLA molecules affect susceptibility to many diseases, most of which are immunologically mediated. In almost all instances, the mechanism that accounts for that predisposition is not known. The microbiome has now been implicated in a long list of diseases, many of which are immunologically mediated. Our studies suggest that HLA molecules could be important factors that contribute to the heterogeneity of the microbiome and operate at least partially through this mechanism in the pathogenesis of many different diseases, not just AS and RA. Consistent with this hypothesis, HLA-microbiome associations have been described in reactive arthritis (62), IBD (63), celiac disease (64) and in healthy individuals (24, 65).

The hypothesized metabolic changes imbued by dysbiosis in our current work are of interest in light of a recent study by our group in the *HLA-B27* transgenic rat model of spondyloarthritis (66). We observe a number of *HLA-B27* dependent metabolic changes in this model that include enrichment of bile acid metabolism, lysine metabolism, fatty acid metabolism and tryptophan metabolism. All of these pathways were predicted to be enriched in *HLA-B27* positive individuals in our current study (Supplementary Table 4). Importantly, *HLA-B27*-dependent dysbiosis can be observed prior to the onset of disease in this model. Thus, our human and rat studies support the hypothesis that *HLA-B27* dependent dysbiosis is a preceding event in AS pathogenesis and may not merely be secondary to disease.

In conclusion, this study demonstrates that *HLA-B27* and RA-associated *HLA-DRB1* allele carriage in humans influences the gut microbiome. In association with the replicated demonstration of intestinal changes in microbiome in AS, this is consistent with disease models in which HLA molecules interact with the gut microbiome to cause disease. Different models as to how this may occur include effects of *HLA-B27* to favour a more inflammatory gut microbiome, increased invasiveness of the gut mucosa in *HLA-B27* carriers, and/or aberrant immunological responses to bacteria in *HLA-B27* carriers. Similar hypotheses may explain the role of *HLA-DRB1* in driving the immunopathogenesis of RA. Whichever of these models is correct, the data presented here support further research in this field, including into whether manipulation of the gut microbiome may be therapeutic in AS or RA, or even potentially capable of preventing disease in at risk subjects.

## ACKNOWLEDGEMENTS

MAB is funded by a National Health and Medical Research Council (Australia) Senior Principal Research Fellowship. This work was funded by a National Health and Medical Research Council (Australia) project grant (reference APP1065509). MA and JTR receive funding from the Spondylitis Association of America and the Rheumatology Research Foundation and from NIH Grant RO1 EY029266. TDS is a National Institute for Health Research (NIHR) senior investigator. TwinsUK is funded by the Wellcome Trust, Medical Research Council, CDRF, Arthritis Research UK, European Union, the NIHR – funded Clinical Research Facility and Biomedical Research Centre based at Guy’s, and St. Thomas’ NHS Foundation Trust in partnership with King’s College London (Grant code WT081878MA). JTR receives funding from the William and Mary Bauman Foundation, the Stan and Madelle Family Trust, and Research to Prevent Blindness. This study was funded in part by the OHSU Foundation (Collins Medical Trust) to JTR. LK receives funding supported by the Eunice Kennedy Shriver National Institute Of Child Health & Human Development of the National Institutes of Health under Award Number K12HD043488. We wish to acknowledge Dr. Erica Roberson, MD for her valuable contribution to study design; Patrick Stauffer and Sean Davin for technical assistance; and all OHSU patients and staff whom helped contribute samples for this study.

