## Supplementary Information for "HLA alleles associated with risk of ankylosing spondylitis and rheumatoid arthritis influence the gut microbiome"

**Supplementary Figure 1:** PCA comparing the microbiome composition at various sample sites. sPLSDA visualization of these results are shown in Figure 1.

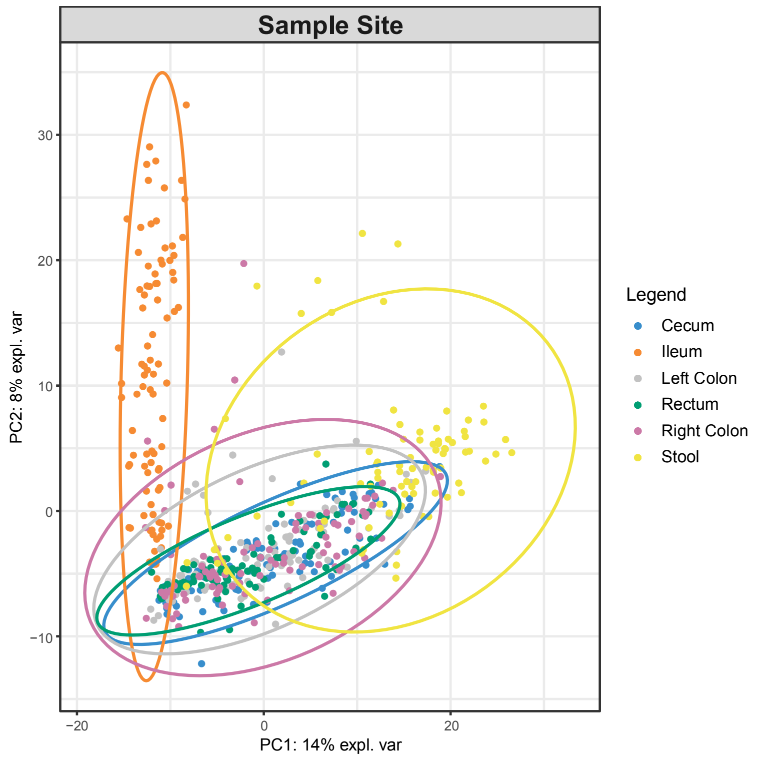


**Supplementary Figure 2: A.** sPLSDA of stool and ileal mucosal microbiome showing minimal overlap between the two (P<0.0001). **B.** PCA of stool and ileal mucosal microbiome.

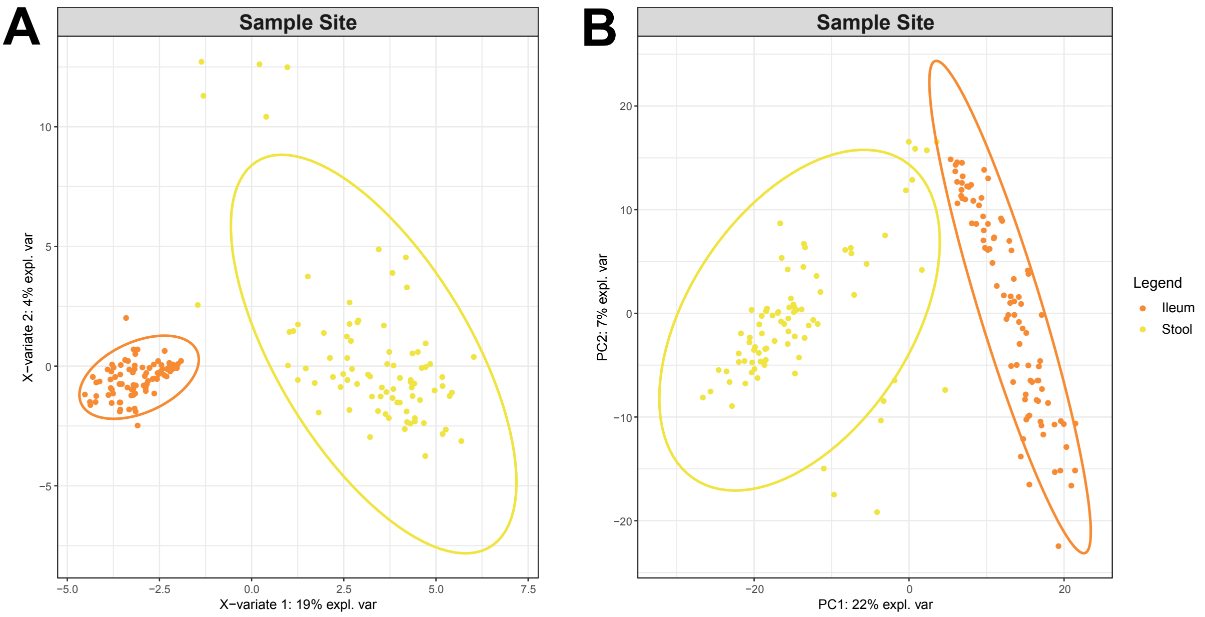

**Supplementary Figure 3: A.** sPLSDA of BMI categories, showing distinct clustering of underweight category compared to overweight, obese and normal. Considering all samples, significant association of BMI with microbiome is observed (P<0.0001). **B.** sPLSDA of BMI categories, having removed the underweight category. **C.** PCA of microbiome composition according to BMI.

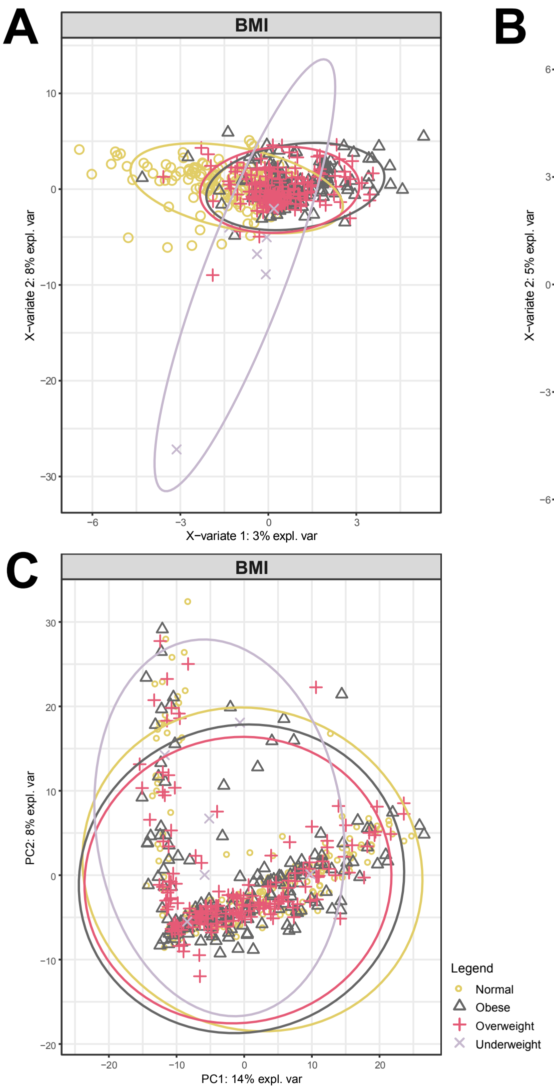

**Supplementary Figure 4: A.** sPLSDA comparing the microbiome composition of female and male genders across all sampling sites, showing substantial overlap but also significant differences between the two (considering all sites, P=0.0004). **B.** PCA of microbiome composition according to gender.

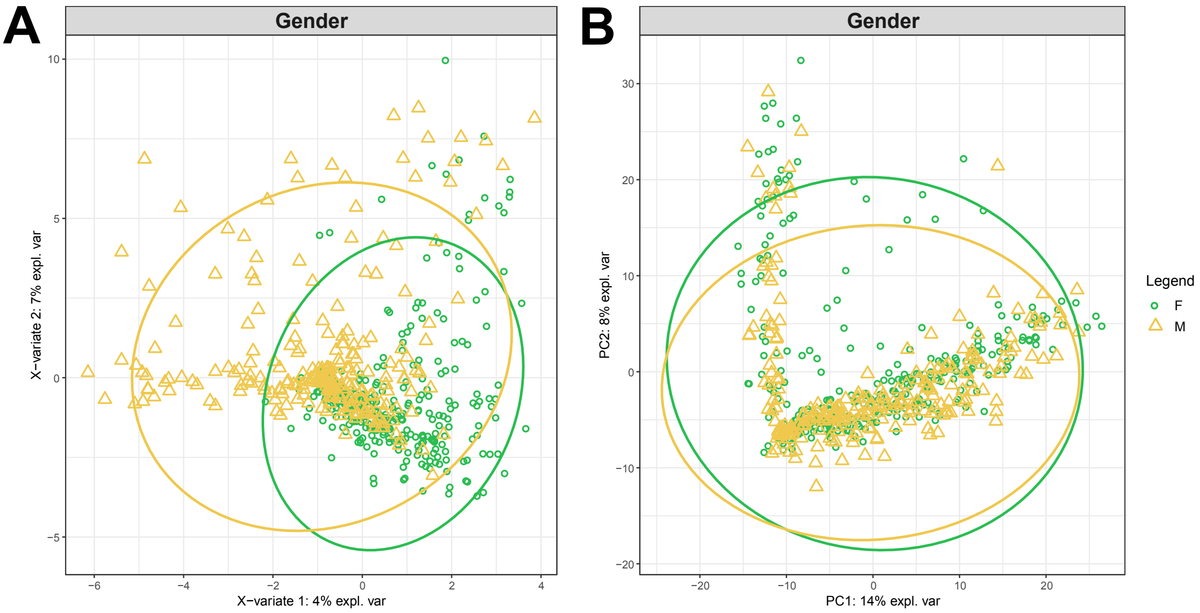

**Supplementary Figure 5:** PCA comparing microbiome composition according to **A.** *HLA-B27* status, **B.** *HLA-DRB1* RA-risk status, and **C.** *HLA-B27* and *HLA-DRB1* RA-risk status in the TwinsUK cohort. sPLSDA visualization of these results are presented in Figure 3.

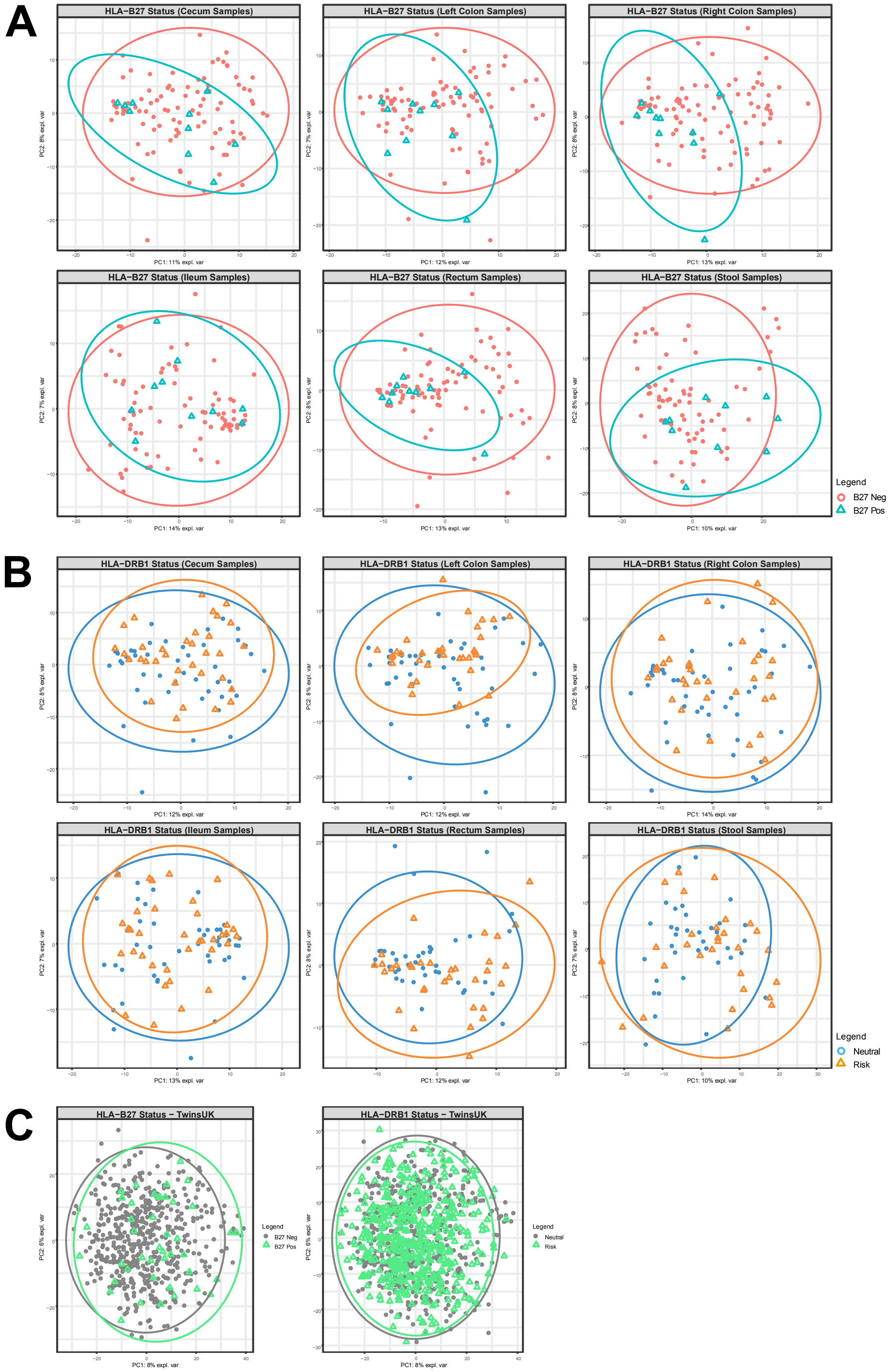

**Supplementary Table 1:** Distribution of *HLA-DRB1* subtypes. Subjects containing *HLA-DRB1* risk and protective alleles were excluded from the analysis. Three of the subjects contained *HLA-DRB1* risk alleles on both haplotypes.

| | Subtype | Count |
|---|---|---|
| **HLA-DRB1 risk** | ***0401** | 16 |
| | ***0404** | 7 |
| | ***0101** | 16 |
| | ***0405** | 0 |
| | ***1402** | 0 |
| **HLA-DRB1 protective** | ***0103** | 1 |
| | ***0402** | 1 |
| | ***0802** | 0 |
| | ***1302** | 6 |
| **HLA-DRB1 neutral** | **Others** | 48 |

**Supplementary Table 2:** Indicator species analysis according to *HLA-B27* status across each sampling site. Associations are color-coded in the following manner: occurred in six sites (red), occurred in five sites (green), occurred in four sites (yellow), occurred in three sites (blue), and occurred in two sites (orange).

| Taxa | HLA Type | Coefficient | P-value | Q-value | Site |
|---|---|---|---|---|---|
| k__Bacteria_p__Actinobacteria_c__Actinobacteria_o__Actinomycetales_f__Actinomycetaceae_g__Varibaculum | B27 Pos | 0.0204 | 0.0047 | 0.5118 | Rectum |
| k__Bacteria_p__Actinobacteria_c__Actinobacteria_o__Actinomycetales_f__Actinomycetaceae_g__Varibaculum_s__ | B27 Pos | 0.0204 | 0.0047 | 0.5118 | Rectum |
| k__Bacteria_p__Bacteroidetes_c__Bacteroidia | B27 Pos | -0.1439 | 0.0330 | 0.6515 | Rectum |
| k__Bacteria_p__Bacteroidetes_c__Bacteroidia_o__Bacteroidales | B27 Pos | -0.1439 | 0.0330 | 0.6515 | Rectum |
| k__Bacteria_p__Bacteroidetes_c__Bacteroidia_o__Bacteroidales_f___Odoribacteraceae__g__Butyricimonas | B27 Pos | -0.0200 | 0.0476 | 0.9111 | Stool |
| k__Bacteria_p__Bacteroidetes_c__Bacteroidia_o__Bacteroidales_f___Odoribacteraceae__g__Butyricimonas_s__ | B27 Pos | -0.0200 | 0.0476 | 0.9111 | Stool |
| k__Bacteria_p__Bacteroidetes_c__Bacteroidia_o__Bacteroidales_f___Paraprevotellaceae_ | B27 Pos | -0.0153 | 0.0478 | 0.7408 | Rectum |
| k__Bacteria_p__Bacteroidetes_c__Bacteroidia_o__Bacteroidales_f__Bacteroidaceae_g__Bacteroides_s__ | B27 Pos | 0.1368 | 0.0367 | 0.9111 | Stool |
| k__Bacteria_p__Bacteroidetes_c__Bacteroidia_o__Bacteroidales_f__Bacteroidaceae_g__Bacteroides_s__caccae | B27 Pos | 0.0130 | 0.0433 | 0.9111 | Stool |
| k__Bacteria_p__Bacteroidetes_c__Bacteroidia_o__Bacteroidales_f__Bacteroidaceae_g__Bacteroides_s__ovatus | B27 Pos | -0.0353 | 0.0139 | 0.8397 | RC |
| k__Bacteria_p__Bacteroidetes_c__Bacteroidia_o__Bacteroidales_f__Bacteroidaceae_g__Bacteroides_s__ovatus | B27 Pos | -0.0411 | 0.0162 | 0.6544 | Cecum |
| k__Bacteria_p__Bacteroidetes_c__Bacteroidia_o__Bacteroidales_f__Bacteroidaceae_g__Bacteroides_s__ovatus | B27 Pos | -0.0408 | 0.0386 | 0.9111 | Stool |
| k__Bacteria_p__Bacteroidetes_c__Bacteroidia_o__Bacteroidales_f__Bacteroidaceae_g__Bacteroides_s__ovatus | B27 Pos | -0.0189 | 0.0448 | 0.7941 | Ileum |
| k__Bacteria_p__Bacteroidetes_c__Bacteroidia_o__Bacteroidales_f__Bacteroidaceae_g__Bacteroides_s__ovatus | B27 Pos | -0.0373 | 0.0493 | 0.7671 | LC |
| k__Bacteria_p__Bacteroidetes_c__Bacteroidia_o__Bacteroidales_f__Prevotellaceae_g__Prevotella_s__ | B27 Pos | 0.0236 | 0.0483 | 0.7941 | Ileum |
| k__Bacteria_p__Bacteroidetes_c__Bacteroidia_o__Bacteroidales_f__Prevotellaceae_g__Prevotella_s__stercorea | B27 Pos | 0.0068 | 0.0458 | 0.7671 | LC |
| k__Bacteria_p__Firmicutes_c__Bacilli_o__Bacillales_f__Staphylococcaceae | B27 Pos | 0.0176 | 0.0045 | 0.3244 | LC |
| k__Bacteria_p__Firmicutes_c__Bacilli_o__Bacillales_f__Staphylococcaceae_g__Staphylococcus | B27 Pos | 0.0176 | 0.0045 | 0.3244 | LC |
| k__Bacteria_p__Firmicutes_c__Bacilli_o__Bacillales_f__Staphylococcaceae_g__Staphylococcus_s__ | B27 Pos | 0.0176 | 0.0045 | 0.3244 | LC |
| k__Bacteria_p__Firmicutes_c__Clostridia_o__Clostridiales_f__Clostridiaceae_g__ | B27 Pos | -0.0170 | 0.0272 | 0.7850 | Cecum |
| k__Bacteria_p__Firmicutes_c__Clostridia_o__Clostridiales_f__Clostridiaceae_g___s__ | B27 Pos | -0.0170 | 0.0272 | 0.7850 | Cecum |
| k__Bacteria_p__Firmicutes_c__Clostridia_o__Clostridiales_f__Lachnospiraceae_g__Blautia_s__obeum | B27 Pos | -0.0414 | 0.0022 | 0.4543 | RC |
| k__Bacteria_p__Firmicutes_c__Clostridia_o__Clostridiales_f__Lachnospiraceae_g__Blautia_s__obeum | B27 Pos | -0.0296 | 0.0305 | 0.7671 | LC |
| k__Bacteria_p__Firmicutes_c__Clostridia_o__Clostridiales_f__Lachnospiraceae_g__Dorea_s__formicigenerans | B27 Pos | -0.0371 | 0.0272 | 0.6515 | Rectum |
| k__Bacteria_p__Firmicutes_c__Clostridia_o__Clostridiales_f__Lachnospiraceae_g__Dorea_s__formicigenerans | B27 Pos | -0.0177 | 0.0461 | 0.9111 | Stool |
| k__Bacteria_p__Firmicutes_c__Clostridia_o__Clostridiales_f__Lachnospiraceae_g__Roseburia | B27 Pos | 0.0516 | 0.0034 | 0.3483 | Stool |
| k__Bacteria_p__Firmicutes_c__Clostridia_o__Clostridiales_f__Lachnospiraceae_g__Roseburia | B27 Pos | 0.0332 | 0.0165 | 0.7044 | LC |
| k__Bacteria_p__Firmicutes_c__Clostridia_o__Clostridiales_f__Lachnospiraceae_g__Roseburia | B27 Pos | 0.0291 | 0.0247 | 0.8397 | RC |
| k__Bacteria_p__Firmicutes_c__Clostridia_o__Clostridiales_f__Lachnospiraceae_g__Roseburia | B27 Pos | 0.0273 | 0.0360 | 0.6515 | Rectum |

| | | | | | |
|---|---|---|---|---|---|
| k__Bacteria_p__Firmicutes_c__Clostridia_o__Clostridiales_f__Lachnospiraceae_g__Roseburia_s__ | B27 Pos | 0.0516 | 0.0034 | 0.3483 | Stool |
| k__Bacteria_p__Firmicutes_c__Clostridia_o__Clostridiales_f__Lachnospiraceae_g__Roseburia_s__ | B27 Pos | 0.0332 | 0.0165 | 0.7044 | LC |
| k__Bacteria_p__Firmicutes_c__Clostridia_o__Clostridiales_f__Lachnospiraceae_g__Roseburia_s__ | B27 Pos | 0.0291 | 0.0247 | 0.8397 | RC |
| k__Bacteria_p__Firmicutes_c__Clostridia_o__Clostridiales_f__Lachnospiraceae_g__Roseburia_s__ | B27 Pos | 0.0273 | 0.0360 | 0.6515 | Rectum |
| k__Bacteria_p__Firmicutes_c__Clostridia_o__Clostridiales_f__Ruminococcaceae_g__Oscillospira | B27 Pos | 0.0170 | 0.0479 | 0.9111 | Stool |
| k__Bacteria_p__Firmicutes_c__Clostridia_o__Clostridiales_f__Ruminococcaceae_g__Oscillospira_s__ | B27 Pos | 0.0170 | 0.0479 | 0.9111 | Stool |
| k__Bacteria_p__Firmicutes_c__Clostridia_o__Clostridiales_f__Veillonellaceae_g__Veillonella_s__dispar | B27 Pos | -0.0202 | 0.0340 | 0.8397 | RC |
| k__Bacteria_p__Proteobacteria_c__Betaproteobacteria_o__Neisseriales | B27 Pos | 0.0306 | 0.0091 | 0.4614 | Cecum |
| k__Bacteria_p__Proteobacteria_c__Betaproteobacteria_o__Neisseriales | B27 Pos | 0.0185 | 0.0093 | 0.4224 | Ileum |
| k__Bacteria_p__Proteobacteria_c__Betaproteobacteria_o__Neisseriales_f__Neisseriaceae | B27 Pos | 0.0306 | 0.0091 | 0.4614 | Cecum |
| k__Bacteria_p__Proteobacteria_c__Betaproteobacteria_o__Neisseriales_f__Neisseriaceae | B27 Pos | 0.0185 | 0.0093 | 0.4224 | Ileum |
| k__Bacteria_p__Proteobacteria_c__Betaproteobacteria_o__Neisseriales_f__Neisseriaceae_g__ | B27 Pos | 0.0185 | 0.0093 | 0.4224 | Ileum |
| k__Bacteria_p__Proteobacteria_c__Betaproteobacteria_o__Neisseriales_f__Neisseriaceae_g___s__ | B27 Pos | 0.0185 | 0.0093 | 0.4224 | Ileum |
| k__Bacteria_p__Proteobacteria_c__Betaproteobacteria_o__Neisseriales_f__Neisseriaceae_g__Neisseria | B27 Pos | 0.0306 | 0.0091 | 0.4614 | Cecum |
| k__Bacteria_p__Proteobacteria_c__Betaproteobacteria_o__Neisseriales_f__Neisseriaceae_g__Neisseria_s__cinerea | B27 Pos | 0.0306 | 0.0091 | 0.4614 | Cecum |
| k__Bacteria_p__Synergistetes_c__Synergistia | B27 Pos | 0.0247 | 0.0263 | 0.6515 | Rectum |
| k__Bacteria_p__Synergistetes_c__Synergistia_o__Synergistales | B27 Pos | 0.0247 | 0.0263 | 0.6515 | Rectum |
| k__Bacteria_p__Synergistetes_c__Synergistia_o__Synergistales_f__ | B27 Pos | 0.0247 | 0.0263 | 0.6515 | Rectum |
| k__Bacteria_p__Synergistetes_c__Synergistia_o__Synergistales_f___g__ | B27 Pos | 0.0247 | 0.0263 | 0.6515 | Rectum |
| k__Bacteria_p__Synergistetes_c__Synergistia_o__Synergistales_f___g___s__ | B27 Pos | 0.0247 | 0.0263 | 0.6515 | Rectum |

| **Taxa** | **Number of tissue sites** |
|---|---|
| k__Bacteria_p__Bacteroidetes_c__Bacteroidia_o__Bacteroidales_f__Bacteroidaceae_g__Bacteroides_s__ovatus | 5 |
| k__Bacteria_p__Firmicutes_c__Clostridia_o__Clostridiales_f__Lachnospiraceae_g__Roseburia | 4 |
| k__Bacteria_p__Firmicutes_c__Clostridia_o__Clostridiales_f__Lachnospiraceae_g__Roseburia_s__ | 4 |
| k__Bacteria_p__Firmicutes_c__Clostridia_o__Clostridiales_f__Lachnospiraceae_g__Blautia_s__obeum | 2 |
| k__Bacteria_p__Firmicutes_c__Clostridia_o__Clostridiales_f__Lachnospiraceae_g__Dorea_s__formicigenerans | 2 |
| k__Bacteria_p__Proteobacteria_c__Betaproteobacteria_o__Neisseriales | 2 |
| k__Bacteria_p__Proteobacteria_c__Betaproteobacteria_o__Neisseriales_f__Neisseriaceae | 2 |
| k__Bacteria_p__Actinobacteria_c__Actinobacteria_o__Actinomycetales_f__Actinomycetaceae_g__Varibaculum | 1 |
| k__Bacteria_p__Actinobacteria_c__Actinobacteria_o__Actinomycetales_f__Actinomycetaceae_g__Varibaculum_s__ | 1 |
| k__Bacteria_p__Bacteroidetes_c__Bacteroidia | 1 |

| | |
|---|---|
| k__Bacteria_p__Bacteroidetes_c__Bacteroidia_o__Bacteroidales | 1 |
| k__Bacteria_p__Bacteroidetes_c__Bacteroidia_o__Bacteroidales_f__Bacteroidaceae_g__Bacteroides_s__ | 1 |
| k__Bacteria_p__Bacteroidetes_c__Bacteroidia_o__Bacteroidales_f__Bacteroidaceae_g__Bacteroides_s__caccae | 1 |
| k__Bacteria_p__Bacteroidetes_c__Bacteroidia_o__Bacteroidales_f___Odoribacteraceae__g__Butyricimonas | 1 |
| k__Bacteria_p__Bacteroidetes_c__Bacteroidia_o__Bacteroidales_f___Odoribacteraceae__g__Butyricimonas_s__ | 1 |
| k__Bacteria_p__Bacteroidetes_c__Bacteroidia_o__Bacteroidales_f___Paraprevotellaceae_ | 1 |
| k__Bacteria_p__Bacteroidetes_c__Bacteroidia_o__Bacteroidales_f__Prevotellaceae_g__Prevotella_s__ | 1 |
| k__Bacteria_p__Bacteroidetes_c__Bacteroidia_o__Bacteroidales_f__Prevotellaceae_g__Prevotella_s__stercorea | 1 |
| k__Bacteria_p__Firmicutes_c__Bacilli_o__Bacillales_f__Staphylococcaceae | 1 |
| k__Bacteria_p__Firmicutes_c__Bacilli_o__Bacillales_f__Staphylococcaceae_g__Staphylococcus | 1 |
| k__Bacteria_p__Firmicutes_c__Bacilli_o__Bacillales_f__Staphylococcaceae_g__Staphylococcus_s__ | 1 |
| k__Bacteria_p__Firmicutes_c__Clostridia_o__Clostridiales_f__Clostridiaceae_g__ | 1 |
| k__Bacteria_p__Firmicutes_c__Clostridia_o__Clostridiales_f__Clostridiaceae_g___s__ | 1 |
| k__Bacteria_p__Firmicutes_c__Clostridia_o__Clostridiales_f__Ruminococcaceae_g__Oscillospira | 1 |
| k__Bacteria_p__Firmicutes_c__Clostridia_o__Clostridiales_f__Ruminococcaceae_g__Oscillospira_s__ | 1 |
| k__Bacteria_p__Firmicutes_c__Clostridia_o__Clostridiales_f__Veillonellaceae_g__Veillonella_s__dispar | 1 |
| k__Bacteria_p__Proteobacteria_c__Betaproteobacteria_o__Neisseriales_f__Neisseriaceae_g__ | 1 |
| k__Bacteria_p__Proteobacteria_c__Betaproteobacteria_o__Neisseriales_f__Neisseriaceae_g__Neisseria | 1 |
| k__Bacteria_p__Proteobacteria_c__Betaproteobacteria_o__Neisseriales_f__Neisseriaceae_g__Neisseria_s__cinerea | 1 |
| k__Bacteria_p__Proteobacteria_c__Betaproteobacteria_o__Neisseriales_f__Neisseriaceae_g___s__ | 1 |
| k__Bacteria_p__Synergistetes_c__Synergistia | 1 |
| k__Bacteria_p__Synergistetes_c__Synergistia_o__Synergistales | 1 |
| k__Bacteria_p__Synergistetes_c__Synergistia_o__Synergistales_f__ | 1 |
| k__Bacteria_p__Synergistetes_c__Synergistia_o__Synergistales_f___g__ | 1 |
| k__Bacteria_p__Synergistetes_c__Synergistia_o__Synergistales_f___g___s__ | 1 |

**Supplementary Table 3:** Indicator species analysis according to *HLA-DRB1* status across each sampling site. Enrichment or depletion is measured relative to the neutral risk genotype. Associations are color-coded in the following manner: occurred in six sites (red), occurred in five sites (green), occurred in four sites (yellow), occurred in three sites (blue), and occurred in two sites (orange).

| Taxa | DRB1 type | Coefficient | P-value | Q-value | Site |
|---|---|---|---|---|---|
| k__Bacteria_p__Actinobacteria_c__Actinobacteria | Protective | 0.0411 | 0.0037 | 0.4823 | Stool |
| k__Bacteria_p__Actinobacteria_c__Actinobacteria | Protective | 0.0198 | 0.0476 | 1.0000 | LC |
| k__Bacteria_p__Actinobacteria_c__Actinobacteria | Risk | 0.0183 | 0.0079 | 0.8664 | RC |
| k__Bacteria_p__Actinobacteria_c__Actinobacteria_o__Actinomycetales_f__Actinomycetaceae | Risk | 0.0036 | 0.0283 | 1.0000 | Cecum |
| k__Bacteria_p__Actinobacteria_c__Actinobacteria_o__Actinomycetales_f__Actinomycetaceae_g__Varibaculum | Protective | 0.0159 | 0.0294 | 1.0000 | Rectum |
| k__Bacteria_p__Actinobacteria_c__Actinobacteria_o__Actinomycetales_f__Actinomycetaceae_g__Varibaculum_s__ | Protective | 0.0159 | 0.0294 | 1.0000 | Rectum |
| k__Bacteria_p__Actinobacteria_c__Actinobacteria_o__Bifidobacteriales | Protective | 0.0448 | 0.0020 | 0.3060 | Stool |
| k__Bacteria_p__Actinobacteria_c__Actinobacteria_o__Bifidobacteriales | Protective | 0.0162 | 0.0278 | 1.0000 | LC |
| k__Bacteria_p__Actinobacteria_c__Actinobacteria_o__Bifidobacteriales | Risk | 0.0099 | 0.0226 | 0.8664 | RC |
| k__Bacteria_p__Actinobacteria_c__Actinobacteria_o__Bifidobacteriales_f__Bifidobacteriaceae | Protective | 0.0448 | 0.0020 | 0.3060 | Stool |
| k__Bacteria_p__Actinobacteria_c__Actinobacteria_o__Bifidobacteriales_f__Bifidobacteriaceae | Protective | 0.0162 | 0.0278 | 1.0000 | LC |
| k__Bacteria_p__Actinobacteria_c__Actinobacteria_o__Bifidobacteriales_f__Bifidobacteriaceae | Risk | 0.0099 | 0.0226 | 0.8664 | RC |
| k__Bacteria_p__Actinobacteria_c__Actinobacteria_o__Bifidobacteriales_f__Bifidobacteriaceae_g__Bifidobacterium | Protective | 0.0495 | 0.0006 | 0.2936 | Stool |
| k__Bacteria_p__Actinobacteria_c__Actinobacteria_o__Bifidobacteriales_f__Bifidobacteriaceae_g__Bifidobacterium | Protective | 0.0165 | 0.0251 | 1.0000 | LC |
| k__Bacteria_p__Actinobacteria_c__Actinobacteria_o__Bifidobacteriales_f__Bifidobacteriaceae_g__Bifidobacterium | Risk | 0.0100 | 0.0212 | 0.8664 | RC |
| k__Bacteria_p__Actinobacteria_c__Actinobacteria_o__Bifidobacteriales_f__Bifidobacteriaceae_g__Bifidobacterium_s__ | Protective | 0.0066 | 0.0096 | 0.9354 | Stool |
| k__Bacteria_p__Actinobacteria_c__Actinobacteria_o__Bifidobacteriales_f__Bifidobacteriaceae_g__Bifidobacterium_s__ | Risk | 0.0169 | 0.0203 | 1.0000 | Rectum |
| k__Bacteria_p__Actinobacteria_c__Actinobacteria_o__Bifidobacteriales_f__Bifidobacteriaceae_g__Bifidobacterium_s__ | Risk | 0.0119 | 0.0299 | 1.0000 | Ileum |
| k__Bacteria_p__Actinobacteria_c__Actinobacteria_o__Bifidobacteriales_f__Bifidobacteriaceae_g__Bifidobacterium_s__adolescentis | Protective | 0.0129 | 0.0098 | 0.8210 | LC |
| k__Bacteria_p__Actinobacteria_c__Actinobacteria_o__Bifidobacteriales_f__Bifidobacteriaceae_g__Bifidobacterium_s__adolescentis | Risk | 0.0066 | 0.0189 | 1.0000 | LC |
| k__Bacteria_p__Actinobacteria_c__Actinobacteria_o__Bifidobacteriales_f__Bifidobacteriaceae_g__Bifidobacterium_s__longum | Risk | 0.0126 | 0.0002 | 0.1689 | RC |
| k__Bacteria_p__Actinobacteria_c__Actinobacteria_o__Bifidobacteriales_f__Bifidobacteriaceae_g__Bifidobacterium_s__longum | Risk | 0.0046 | 0.0070 | 1.0000 | Cecum |
| k__Bacteria_p__Actinobacteria_c__Actinobacteria_o__Bifidobacteriales_f__Bifidobacteriaceae_g__Bifidobacterium_s__longum | Risk | 0.0033 | 0.0217 | 1.0000 | Rectum |
| k__Bacteria_p__Bacteroidetes_c__Bacteroidia_o__Bacteroidales_f___Odoribacteraceae_ | Risk | -0.0240 | 0.0327 | 1.0000 | Cecum |
| k__Bacteria_p__Bacteroidetes_c__Bacteroidia_o__Bacteroidales_f__Bacteroidaceae_g__Bacteroides_s__eggerthii | Risk | -0.0040 | 0.0375 | 1.0000 | Ileum |
| k__Bacteria_p__Bacteroidetes_c__Bacteroidia_o__Bacteroidales_f__Bacteroidaceae_g__Bacteroides_s__uniformis | Risk | -0.0319 | 0.0466 | 1.0000 | Rectum |
| k__Bacteria_p__Bacteroidetes_c__Bacteroidia_o__Bacteroidales_f__Rikenellaceae | Risk | -0.0309 | 0.0105 | 0.8664 | RC |
| k__Bacteria_p__Bacteroidetes_c__Bacteroidia_o__Bacteroidales_f__Rikenellaceae | Risk | -0.0253 | 0.0317 | 1.0000 | Cecum |

| | | | | | |
|---|---|---|---|---|---|
| k__Bacteria_p__Bacteroidetes_c__Bacteroidia_o__Bacteroidales_f__Rikenellaceae_g__ | Risk | -0.0290 | 0.0160 | 0.8664 | RC |
| k__Bacteria_p__Bacteroidetes_c__Bacteroidia_o__Bacteroidales_f__Rikenellaceae_g___s__ | Risk | -0.0290 | 0.0160 | 0.8664 | RC |
| k__Bacteria_p__Firmicutes_c__Bacilli_o__Bacillales_f__Staphylococcaceae | Protective | -0.0151 | 0.0386 | 1.0000 | Cecum |
| k__Bacteria_p__Firmicutes_c__Bacilli_o__Bacillales_f__Staphylococcaceae_g__Staphylococcus | Protective | -0.0151 | 0.0386 | 1.0000 | Cecum |
| k__Bacteria_p__Firmicutes_c__Bacilli_o__Bacillales_f__Staphylococcaceae_g__Staphylococcus_s__ | Protective | -0.0151 | 0.0386 | 1.0000 | Cecum |
| k__Bacteria_p__Firmicutes_c__Bacilli_o__Lactobacillales_f__Lactobacillaceae | Protective | 0.0409 | 0.0075 | 0.7192 | Rectum |
| k__Bacteria_p__Firmicutes_c__Bacilli_o__Lactobacillales_f__Lactobacillaceae_g__Lactobacillus | Protective | 0.0409 | 0.0075 | 0.7192 | Rectum |
| k__Bacteria_p__Firmicutes_c__Bacilli_o__Lactobacillales_f__Lactobacillaceae_g__Lactobacillus_s__ | Protective | 0.0409 | 0.0075 | 0.7192 | Rectum |
| k__Bacteria_p__Firmicutes_c__Bacilli_o__Turicibacterales | Protective | 0.0043 | 0.0227 | 1.0000 | Stool |
| k__Bacteria_p__Firmicutes_c__Bacilli_o__Turicibacterales_f__Turicibacteraceae | Protective | 0.0043 | 0.0227 | 1.0000 | Stool |
| k__Bacteria_p__Firmicutes_c__Bacilli_o__Turicibacterales_f__Turicibacteraceae_g__Turicibacter | Protective | 0.0043 | 0.0227 | 1.0000 | Stool |
| k__Bacteria_p__Firmicutes_c__Bacilli_o__Turicibacterales_f__Turicibacteraceae_g__Turicibacter_s__ | Protective | 0.0043 | 0.0227 | 1.0000 | Stool |
| k__Bacteria_p__Firmicutes_c__Clostridia_o__Clostridiales_f___Mogibacteriaceae_ | Protective | 0.0060 | 0.0223 | 1.0000 | Stool |
| k__Bacteria_p__Firmicutes_c__Clostridia_o__Clostridiales_f___Mogibacteriaceae__g__ | Protective | 0.0061 | 0.0184 | 1.0000 | Stool |
| k__Bacteria_p__Firmicutes_c__Clostridia_o__Clostridiales_f___Mogibacteriaceae__g___s__ | Protective | 0.0061 | 0.0184 | 1.0000 | Stool |
| k__Bacteria_p__Firmicutes_c__Clostridia_o__Clostridiales_f___Tissierellaceae__g__Peptoniphilus | Risk | -0.0186 | 0.0130 | 1.0000 | Rectum |
| k__Bacteria_p__Firmicutes_c__Clostridia_o__Clostridiales_f___Tissierellaceae__g__Peptoniphilus_s__ | Risk | -0.0186 | 0.0130 | 1.0000 | Rectum |
| k__Bacteria_p__Firmicutes_c__Clostridia_o__Clostridiales_f___Tissierellaceae__g__WAL_1855D | Protective | 0.0100 | 0.0457 | 1.0000 | LC |
| k__Bacteria_p__Firmicutes_c__Clostridia_o__Clostridiales_f___Tissierellaceae__g__WAL_1855D_s__ | Protective | 0.0100 | 0.0457 | 1.0000 | LC |
| k__Bacteria_p__Firmicutes_c__Clostridia_o__Clostridiales_f__Clostridiaceae | Risk | 0.0283 | 0.0063 | 0.7026 | Stool |
| k__Bacteria_p__Firmicutes_c__Clostridia_o__Clostridiales_f__Clostridiaceae | Risk | 0.0214 | 0.0140 | 0.8508 | LC |
| k__Bacteria_p__Firmicutes_c__Clostridia_o__Clostridiales_f__Clostridiaceae | Risk | 0.0221 | 0.0157 | 1.0000 | Rectum |
| k__Bacteria_p__Firmicutes_c__Clostridia_o__Clostridiales_f__Clostridiaceae_g__ | Risk | 0.0158 | 0.0184 | 0.8664 | RC |
| k__Bacteria_p__Firmicutes_c__Clostridia_o__Clostridiales_f__Clostridiaceae_g__ | Risk | 0.0128 | 0.0268 | 1.0000 | Rectum |
| k__Bacteria_p__Firmicutes_c__Clostridia_o__Clostridiales_f__Clostridiaceae_g__ | Risk | 0.0147 | 0.0297 | 1.0000 | LC |
| k__Bacteria_p__Firmicutes_c__Clostridia_o__Clostridiales_f__Clostridiaceae_g__ | Risk | 0.0120 | 0.0447 | 1.0000 | Stool |
| k__Bacteria_p__Firmicutes_c__Clostridia_o__Clostridiales_f__Clostridiaceae_g___s__ | Risk | 0.0158 | 0.0184 | 0.8664 | RC |
| k__Bacteria_p__Firmicutes_c__Clostridia_o__Clostridiales_f__Clostridiaceae_g___s__ | Risk | 0.0128 | 0.0268 | 1.0000 | Rectum |
| k__Bacteria_p__Firmicutes_c__Clostridia_o__Clostridiales_f__Clostridiaceae_g___s__ | Risk | 0.0147 | 0.0297 | 1.0000 | LC |
| k__Bacteria_p__Firmicutes_c__Clostridia_o__Clostridiales_f__Clostridiaceae_g___s__ | Risk | 0.0120 | 0.0447 | 1.0000 | Stool |
| k__Bacteria_p__Firmicutes_c__Clostridia_o__Clostridiales_f__Clostridiaceae_g__SMB53 | Risk | 0.0123 | 0.0011 | 0.2936 | Stool |
| k__Bacteria_p__Firmicutes_c__Clostridia_o__Clostridiales_f__Clostridiaceae_g__SMB53 | Risk | 0.0062 | 0.0053 | 0.7192 | Rectum |
| k__Bacteria_p__Firmicutes_c__Clostridia_o__Clostridiales_f__Clostridiaceae_g__SMB53 | Risk | 0.0047 | 0.0415 | 1.0000 | RC |

| | | | | | |
|---|---|---|---|---|---|
| k__Bacteria_p__Firmicutes_c__Clostridia_o__Clostridiales_f__Clostridiaceae_g__SMB53_s__ | Risk | 0.0123 | 0.0011 | 0.2936 | Stool |
| k__Bacteria_p__Firmicutes_c__Clostridia_o__Clostridiales_f__Clostridiaceae_g__SMB53_s__ | Risk | 0.0062 | 0.0053 | 0.7192 | Rectum |
| k__Bacteria_p__Firmicutes_c__Clostridia_o__Clostridiales_f__Clostridiaceae_g__SMB53_s__ | Risk | 0.0047 | 0.0415 | 1.0000 | RC |
| k__Bacteria_p__Firmicutes_c__Clostridia_o__Clostridiales_f__Lachnospiraceae | Risk | 0.0816 | 0.0058 | 0.8210 | LC |
| k__Bacteria_p__Firmicutes_c__Clostridia_o__Clostridiales_f__Lachnospiraceae | Risk | 0.0660 | 0.0305 | 1.0000 | RC |
| k__Bacteria_p__Firmicutes_c__Clostridia_o__Clostridiales_f__Lachnospiraceae | Risk | 0.0569 | 0.0312 | 1.0000 | Rectum |
| k__Bacteria_p__Firmicutes_c__Clostridia_o__Clostridiales_f__Lachnospiraceae | Risk | 0.0651 | 0.0339 | 1.0000 | Ileum |
| k__Bacteria_p__Firmicutes_c__Clostridia_o__Clostridiales_f__Lachnospiraceae_g__ | Risk | 0.0688 | 0.0074 | 0.8210 | LC |
| k__Bacteria_p__Firmicutes_c__Clostridia_o__Clostridiales_f__Lachnospiraceae_g__ | Risk | 0.0644 | 0.0091 | 1.0000 | Cecum |
| k__Bacteria_p__Firmicutes_c__Clostridia_o__Clostridiales_f__Lachnospiraceae_g__ | Risk | 0.0513 | 0.0176 | 1.0000 | Rectum |
| k__Bacteria_p__Firmicutes_c__Clostridia_o__Clostridiales_f__Lachnospiraceae_g__ | Risk | 0.0590 | 0.0219 | 0.9897 | Ileum |
| k__Bacteria_p__Firmicutes_c__Clostridia_o__Clostridiales_f__Lachnospiraceae_g__ | Risk | 0.0558 | 0.0329 | 1.0000 | RC |
| k__Bacteria_p__Firmicutes_c__Clostridia_o__Clostridiales_f__Lachnospiraceae_g___Ruminococcus__s__gnavus | Risk | 0.0152 | 0.0183 | 0.9897 | Ileum |
| k__Bacteria_p__Firmicutes_c__Clostridia_o__Clostridiales_f__Lachnospiraceae_g___s__ | Risk | 0.0688 | 0.0074 | 0.8210 | LC |
| k__Bacteria_p__Firmicutes_c__Clostridia_o__Clostridiales_f__Lachnospiraceae_g___s__ | Risk | 0.0644 | 0.0091 | 1.0000 | Cecum |
| k__Bacteria_p__Firmicutes_c__Clostridia_o__Clostridiales_f__Lachnospiraceae_g___s__ | Risk | 0.0513 | 0.0176 | 1.0000 | Rectum |
| k__Bacteria_p__Firmicutes_c__Clostridia_o__Clostridiales_f__Lachnospiraceae_g___s__ | Risk | 0.0590 | 0.0219 | 0.9897 | Ileum |
| k__Bacteria_p__Firmicutes_c__Clostridia_o__Clostridiales_f__Lachnospiraceae_g___s__ | Risk | 0.0558 | 0.0329 | 1.0000 | RC |
| k__Bacteria_p__Firmicutes_c__Clostridia_o__Clostridiales_f__Lachnospiraceae_g__Blautia_s__obeum | Risk | 0.0192 | 0.0352 | 1.0000 | LC |
| k__Bacteria_p__Firmicutes_c__Clostridia_o__Clostridiales_f__Lachnospiraceae_g__Blautia_s__producta | Risk | -0.0121 | 0.0465 | 1.0000 | Stool |
| k__Bacteria_p__Firmicutes_c__Clostridia_o__Clostridiales_f__Lachnospiraceae_g__Coprococcus | Risk | 0.0255 | 0.0072 | 0.7192 | Rectum |
| k__Bacteria_p__Firmicutes_c__Clostridia_o__Clostridiales_f__Lachnospiraceae_g__Coprococcus | Risk | 0.0215 | 0.0306 | 1.0000 | LC |
| k__Bacteria_p__Firmicutes_c__Clostridia_o__Clostridiales_f__Lachnospiraceae_g__Coprococcus_s__ | Risk | 0.0253 | 0.0073 | 0.7192 | Rectum |
| k__Bacteria_p__Firmicutes_c__Clostridia_o__Clostridiales_f__Lachnospiraceae_g__Coprococcus_s__ | Risk | 0.0219 | 0.0246 | 1.0000 | LC |
| k__Bacteria_p__Firmicutes_c__Clostridia_o__Clostridiales_f__Lachnospiraceae_g__Coprococcus_s__eutactus | Protective | 0.0286 | 0.0310 | 1.0000 | Stool |
| k__Bacteria_p__Firmicutes_c__Clostridia_o__Clostridiales_f__Lachnospiraceae_g__Coprococcus_s__eutactus | Protective | 0.0123 | 0.0443 | 1.0000 | LC |
| k__Bacteria_p__Firmicutes_c__Clostridia_o__Clostridiales_f__Lachnospiraceae_g__Dorea | Protective | 0.0197 | 0.0402 | 1.0000 | Stool |
| k__Bacteria_p__Firmicutes_c__Clostridia_o__Clostridiales_f__Lachnospiraceae_g__Dorea_s__ | Risk | 0.0205 | 0.0231 | 0.8664 | RC |
| k__Bacteria_p__Firmicutes_c__Clostridia_o__Clostridiales_f__Lachnospiraceae_g__Dorea_s__formicigenerans | Risk | 0.0224 | 0.0214 | 0.9897 | Ileum |
| k__Bacteria_p__Firmicutes_c__Clostridia_o__Clostridiales_f__Lachnospiraceae_g__Roseburia | Protective | 0.0292 | 0.0464 | 1.0000 | Rectum |
| k__Bacteria_p__Firmicutes_c__Clostridia_o__Clostridiales_f__Lachnospiraceae_g__Roseburia | Risk | 0.0186 | 0.0278 | 1.0000 | Rectum |
| k__Bacteria_p__Firmicutes_c__Clostridia_o__Clostridiales_f__Lachnospiraceae_g__Roseburia_s__ | Protective | 0.0292 | 0.0464 | 1.0000 | Rectum |
| k__Bacteria_p__Firmicutes_c__Clostridia_o__Clostridiales_f__Lachnospiraceae_g__Roseburia_s__ | Risk | 0.0186 | 0.0278 | 1.0000 | Rectum |

| | | | | | |
|---|---|---|---|---|---|
| k__Bacteria_p__Firmicutes_c__Clostridia_o__Clostridiales_f__Peptococcaceae | Risk | -0.0115 | 0.0318 | 1.0000 | LC |
| k__Bacteria_p__Firmicutes_c__Clostridia_o__Clostridiales_f__Peptococcaceae | Risk | -0.0084 | 0.0390 | 1.0000 | RC |
| k__Bacteria_p__Firmicutes_c__Clostridia_o__Clostridiales_f__Peptococcaceae_g__rc4_4 | Risk | -0.0081 | 0.0319 | 1.0000 | Cecum |
| k__Bacteria_p__Firmicutes_c__Clostridia_o__Clostridiales_f__Peptococcaceae_g__rc4_4 | Risk | -0.0085 | 0.0471 | 1.0000 | LC |
| k__Bacteria_p__Firmicutes_c__Clostridia_o__Clostridiales_f__Peptococcaceae_g__rc4_4_s__ | Risk | -0.0081 | 0.0319 | 1.0000 | Cecum |
| k__Bacteria_p__Firmicutes_c__Clostridia_o__Clostridiales_f__Peptococcaceae_g__rc4_4_s__ | Risk | -0.0085 | 0.0471 | 1.0000 | LC |
| k__Bacteria_p__Firmicutes_c__Clostridia_o__Clostridiales_f__Peptostreptococcaceae | Risk | 0.0081 | 0.0380 | 1.0000 | Stool |
| k__Bacteria_p__Firmicutes_c__Clostridia_o__Clostridiales_f__Peptostreptococcaceae_g__ | Risk | 0.0081 | 0.0380 | 1.0000 | Stool |
| k__Bacteria_p__Firmicutes_c__Clostridia_o__Clostridiales_f__Peptostreptococcaceae_g___s__ | Risk | 0.0081 | 0.0380 | 1.0000 | Stool |
| k__Bacteria_p__Firmicutes_c__Clostridia_o__Clostridiales_f__Ruminococcaceae | Protective | 0.1013 | 0.0367 | 1.0000 | Ileum |
| k__Bacteria_p__Firmicutes_c__Clostridia_o__Clostridiales_f__Ruminococcaceae_g__Oscillospira | Risk | -0.0163 | 0.0463 | 1.0000 | LC |
| k__Bacteria_p__Firmicutes_c__Clostridia_o__Clostridiales_f__Ruminococcaceae_g__Oscillospira_s__ | Risk | -0.0163 | 0.0463 | 1.0000 | LC |
| k__Bacteria_p__Firmicutes_c__Clostridia_o__Clostridiales_f__Veillonellaceae | Protective | 0.0532 | 0.0153 | 0.8671 | LC |
| k__Bacteria_p__Firmicutes_c__Clostridia_o__Clostridiales_f__Veillonellaceae | Protective | 0.0468 | 0.0225 | 0.8664 | RC |
| k__Bacteria_p__Firmicutes_c__Clostridia_o__Clostridiales_f__Veillonellaceae | Protective | 0.0284 | 0.0347 | 1.0000 | Stool |
| k__Bacteria_p__Firmicutes_c__Clostridia_o__Clostridiales_f__Veillonellaceae_g__Acidaminococcus | Risk | 0.0107 | 0.0458 | 1.0000 | Stool |
| k__Bacteria_p__Firmicutes_c__Clostridia_o__Clostridiales_f__Veillonellaceae_g__Acidaminococcus_s__ | Risk | 0.0107 | 0.0458 | 1.0000 | Stool |
| k__Bacteria_p__Firmicutes_c__Clostridia_o__Clostridiales_f__Veillonellaceae_g__Dialister | Protective | 0.0326 | 0.0204 | 1.0000 | Rectum |
| k__Bacteria_p__Firmicutes_c__Clostridia_o__Clostridiales_f__Veillonellaceae_g__Dialister | Risk | 0.0103 | 0.0179 | 1.0000 | Stool |
| k__Bacteria_p__Firmicutes_c__Clostridia_o__Clostridiales_f__Veillonellaceae_g__Dialister_s__ | Protective | 0.0326 | 0.0204 | 1.0000 | Rectum |
| k__Bacteria_p__Firmicutes_c__Clostridia_o__Clostridiales_f__Veillonellaceae_g__Dialister_s__ | Risk | 0.0103 | 0.0179 | 1.0000 | Stool |
| k__Bacteria_p__Firmicutes_c__Clostridia_o__Clostridiales_f__Veillonellaceae_g__Phascolarctobacterium | Risk | -0.0204 | 0.0483 | 1.0000 | Ileum |
| k__Bacteria_p__Firmicutes_c__Clostridia_o__Clostridiales_f__Veillonellaceae_g__Phascolarctobacterium_s__ | Risk | -0.0204 | 0.0483 | 1.0000 | Ileum |
| k__Bacteria_p__Firmicutes_c__Clostridia_o__Clostridiales_f__Veillonellaceae_g__Veillonella | Risk | 0.0154 | 0.0289 | 1.0000 | RC |
| k__Bacteria_p__Firmicutes_c__Clostridia_o__Clostridiales_f__Veillonellaceae_g__Veillonella_s__dispar | Risk | 0.0162 | 0.0099 | 0.8664 | RC |
| k__Bacteria_p__Firmicutes_c__Erysipelotrichi | Risk | 0.0043 | 0.0086 | 0.7276 | Ileum |
| k__Bacteria_p__Firmicutes_c__Erysipelotrichi_o__Erysipelotrichales | Risk | 0.0043 | 0.0086 | 0.7276 | Ileum |
| k__Bacteria_p__Firmicutes_c__Erysipelotrichi_o__Erysipelotrichales_f__Erysipelotrichaceae | Risk | 0.0043 | 0.0086 | 0.7276 | Ileum |
| k__Bacteria_p__Firmicutes_c__Erysipelotrichi_o__Erysipelotrichales_f__Erysipelotrichaceae_g__ | Risk | 0.0043 | 0.0086 | 0.7276 | Ileum |
| k__Bacteria_p__Firmicutes_c__Erysipelotrichi_o__Erysipelotrichales_f__Erysipelotrichaceae_g___s__ | Risk | 0.0043 | 0.0086 | 0.7276 | Ileum |
| k__Bacteria_p__Proteobacteria_c__Alphaproteobacteria | Risk | -0.0180 | 0.0402 | 1.0000 | Rectum |
| k__Bacteria_p__Proteobacteria_c__Alphaproteobacteria_o__RF32 | Risk | -0.0063 | 0.0063 | 0.8210 | LC |
| k__Bacteria_p__Proteobacteria_c__Alphaproteobacteria_o__RF32 | Risk | -0.0039 | 0.0172 | 0.8664 | RC |

| | | | | | |
|---|---|---|---|---|---|
| k__Bacteria_p__Proteobacteria_c__Alphaproteobacteria_o__RF32 | Risk | -0.0180 | 0.0402 | 1.0000 | Rectum |
| k__Bacteria_p__Proteobacteria_c__Alphaproteobacteria_o__RF32_f__ | Risk | -0.0063 | 0.0063 | 0.8210 | LC |
| k__Bacteria_p__Proteobacteria_c__Alphaproteobacteria_o__RF32_f__ | Risk | -0.0039 | 0.0172 | 0.8664 | RC |
| k__Bacteria_p__Proteobacteria_c__Alphaproteobacteria_o__RF32_f__ | Risk | -0.0180 | 0.0402 | 1.0000 | Rectum |
| k__Bacteria_p__Proteobacteria_c__Alphaproteobacteria_o__RF32_f___g__ | Risk | -0.0063 | 0.0063 | 0.8210 | LC |
| k__Bacteria_p__Proteobacteria_c__Alphaproteobacteria_o__RF32_f___g__ | Risk | -0.0039 | 0.0172 | 0.8664 | RC |
| k__Bacteria_p__Proteobacteria_c__Alphaproteobacteria_o__RF32_f___g__ | Risk | -0.0180 | 0.0402 | 1.0000 | Rectum |
| k__Bacteria_p__Proteobacteria_c__Alphaproteobacteria_o__RF32_f___g___s__ | Risk | -0.0063 | 0.0063 | 0.8210 | LC |
| k__Bacteria_p__Proteobacteria_c__Alphaproteobacteria_o__RF32_f___g___s__ | Risk | -0.0039 | 0.0172 | 0.8664 | RC |
| k__Bacteria_p__Proteobacteria_c__Alphaproteobacteria_o__RF32_f___g___s__ | Risk | -0.0180 | 0.0402 | 1.0000 | Rectum |
| k__Bacteria_p__Proteobacteria_c__Deltaproteobacteria | Protective | 0.0272 | 0.0107 | 0.7276 | Ileum |
| k__Bacteria_p__Proteobacteria_c__Deltaproteobacteria | Protective | 0.0334 | 0.0368 | 1.0000 | Rectum |
| k__Bacteria_p__Proteobacteria_c__Deltaproteobacteria | Risk | -0.0173 | 0.0419 | 1.0000 | Cecum |
| k__Bacteria_p__Proteobacteria_c__Deltaproteobacteria_o__Desulfovibrionales | Protective | 0.0272 | 0.0107 | 0.7276 | Ileum |
| k__Bacteria_p__Proteobacteria_c__Deltaproteobacteria_o__Desulfovibrionales | Protective | 0.0334 | 0.0368 | 1.0000 | Rectum |
| k__Bacteria_p__Proteobacteria_c__Deltaproteobacteria_o__Desulfovibrionales | Risk | -0.0173 | 0.0419 | 1.0000 | Cecum |
| k__Bacteria_p__Proteobacteria_c__Deltaproteobacteria_o__Desulfovibrionales_f__Desulfovibrionaceae | Protective | 0.0272 | 0.0107 | 0.7276 | Ileum |
| k__Bacteria_p__Proteobacteria_c__Deltaproteobacteria_o__Desulfovibrionales_f__Desulfovibrionaceae | Protective | 0.0334 | 0.0368 | 1.0000 | Rectum |
| k__Bacteria_p__Proteobacteria_c__Deltaproteobacteria_o__Desulfovibrionales_f__Desulfovibrionaceae | Risk | -0.0173 | 0.0419 | 1.0000 | Cecum |
| k__Bacteria_p__Proteobacteria_c__Deltaproteobacteria_o__Desulfovibrionales_f__Desulfovibrionaceae_g__Bilophila | Protective | 0.0434 | 0.0011 | 0.4636 | Rectum |
| k__Bacteria_p__Proteobacteria_c__Deltaproteobacteria_o__Desulfovibrionales_f__Desulfovibrionaceae_g__Bilophila | Protective | 0.0288 | 0.0030 | 0.7276 | Ileum |
| k__Bacteria_p__Proteobacteria_c__Deltaproteobacteria_o__Desulfovibrionales_f__Desulfovibrionaceae_g__Bilophila | Risk | -0.0172 | 0.0202 | 1.0000 | Cecum |
| k__Bacteria_p__Proteobacteria_c__Deltaproteobacteria_o__Desulfovibrionales_f__Desulfovibrionaceae_g__Bilophila_s__ | Protective | 0.0434 | 0.0011 | 0.4636 | Rectum |
| k__Bacteria_p__Proteobacteria_c__Deltaproteobacteria_o__Desulfovibrionales_f__Desulfovibrionaceae_g__Bilophila_s__ | Protective | 0.0288 | 0.0030 | 0.7276 | Ileum |
| k__Bacteria_p__Proteobacteria_c__Deltaproteobacteria_o__Desulfovibrionales_f__Desulfovibrionaceae_g__Bilophila_s__ | Risk | -0.0172 | 0.0202 | 1.0000 | Cecum |
| k__Bacteria_p__Proteobacteria_c__Deltaproteobacteria_o__Desulfovibrionales_f__Desulfovibrionaceae_g__Desulfovibrio | Protective | 0.0278 | 0.0280 | 1.0000 | LC |
| k__Bacteria_p__Proteobacteria_c__Gammaproteobacteria_o__Enterobacteriales_f__Enterobacteriaceae_g__Klebsiella | Risk | 0.0164 | 0.0121 | 0.7276 | Ileum |
| k__Bacteria_p__Proteobacteria_c__Gammaproteobacteria_o__Enterobacteriales_f__Enterobacteriaceae_g__Klebsiella_s__ | Risk | 0.0164 | 0.0121 | 0.7276 | Ileum |
| k__Bacteria_p__Proteobacteria_c__Gammaproteobacteria_o__Oceanospirillales | Risk | -0.0203 | 0.0420 | 1.0000 | Rectum |
| k__Bacteria_p__Proteobacteria_c__Gammaproteobacteria_o__Oceanospirillales_f__Halomonadaceae | Risk | -0.0203 | 0.0420 | 1.0000 | Rectum |
| k__Bacteria_p__Verrucomicrobia_c__Verrucomicrobiae | Risk | -0.0145 | 0.0015 | 0.2106 | RC |
| k__Bacteria_p__Verrucomicrobia_c__Verrucomicrobiae | Risk | -0.0056 | 0.0126 | 0.8210 | LC |
| k__Bacteria_p__Verrucomicrobia_c__Verrucomicrobiae | Risk | -0.0073 | 0.0247 | 1.0000 | Cecum |

| | | | | | |
|---|---|---|---|---|---|
| k__Bacteria_p__Verrucomicrobia_c__Verrucomicrobiae_o__Verrucomicrobiales | Risk | -0.0145 | 0.0015 | 0.2106 | RC |
| k__Bacteria_p__Verrucomicrobia_c__Verrucomicrobiae_o__Verrucomicrobiales | Risk | -0.0056 | 0.0126 | 0.8210 | LC |
| k__Bacteria_p__Verrucomicrobia_c__Verrucomicrobiae_o__Verrucomicrobiales | Risk | -0.0073 | 0.0247 | 1.0000 | Cecum |
| k__Bacteria_p__Verrucomicrobia_c__Verrucomicrobiae_o__Verrucomicrobiales_f__Verrucomicrobiaceae | Risk | -0.0145 | 0.0015 | 0.2106 | RC |
| k__Bacteria_p__Verrucomicrobia_c__Verrucomicrobiae_o__Verrucomicrobiales_f__Verrucomicrobiaceae | Risk | -0.0056 | 0.0126 | 0.8210 | LC |
| k__Bacteria_p__Verrucomicrobia_c__Verrucomicrobiae_o__Verrucomicrobiales_f__Verrucomicrobiaceae | Risk | -0.0073 | 0.0247 | 1.0000 | Cecum |
| k__Bacteria_p__Verrucomicrobia_c__Verrucomicrobiae_o__Verrucomicrobiales_f__Verrucomicrobiaceae_g__Akkermansia | Risk | -0.0145 | 0.0015 | 0.2106 | RC |
| k__Bacteria_p__Verrucomicrobia_c__Verrucomicrobiae_o__Verrucomicrobiales_f__Verrucomicrobiaceae_g__Akkermansia | Risk | -0.0056 | 0.0126 | 0.8210 | LC |
| k__Bacteria_p__Verrucomicrobia_c__Verrucomicrobiae_o__Verrucomicrobiales_f__Verrucomicrobiaceae_g__Akkermansia | Risk | -0.0073 | 0.0247 | 1.0000 | Cecum |
| k__Bacteria_p__Verrucomicrobia_c__Verrucomicrobiae_o__Verrucomicrobiales_f__Verrucomicrobiaceae_g__Akkermansia_s__muciniphila | Risk | -0.0145 | 0.0015 | 0.2106 | RC |
| k__Bacteria_p__Verrucomicrobia_c__Verrucomicrobiae_o__Verrucomicrobiales_f__Verrucomicrobiaceae_g__Akkermansia_s__muciniphila | Risk | -0.0056 | 0.0126 | 0.8210 | LC |
| k__Bacteria_p__Verrucomicrobia_c__Verrucomicrobiae_o__Verrucomicrobiales_f__Verrucomicrobiaceae_g__Akkermansia_s__muciniphila | Risk | -0.0073 | 0.0247 | 1.0000 | Cecum |

| **Taxa** | **Number of tissue sites** |
|---|---|
| k__Bacteria_p__Firmicutes_c__Clostridia_o__Clostridiales_f__Lachnospiraceae_g__ | 5 |
| k__Bacteria_p__Firmicutes_c__Clostridia_o__Clostridiales_f__Lachnospiraceae_g___s__ | 5 |
| k__Bacteria_p__Firmicutes_c__Clostridia_o__Clostridiales_f__Clostridiaceae_g__ | 4 |
| k__Bacteria_p__Firmicutes_c__Clostridia_o__Clostridiales_f__Clostridiaceae_g___s__ | 4 |
| k__Bacteria_p__Firmicutes_c__Clostridia_o__Clostridiales_f__Lachnospiraceae | 4 |
| k__Bacteria_p__Actinobacteria_c__Actinobacteria | 3 |
| k__Bacteria_p__Actinobacteria_c__Actinobacteria_o__Bifidobacteriales | 3 |
| k__Bacteria_p__Actinobacteria_c__Actinobacteria_o__Bifidobacteriales_f__Bifidobacteriaceae | 3 |
| k__Bacteria_p__Actinobacteria_c__Actinobacteria_o__Bifidobacteriales_f__Bifidobacteriaceae_g__Bifidobacterium | 3 |
| k__Bacteria_p__Actinobacteria_c__Actinobacteria_o__Bifidobacteriales_f__Bifidobacteriaceae_g__Bifidobacterium_s__ | 3 |
| k__Bacteria_p__Actinobacteria_c__Actinobacteria_o__Bifidobacteriales_f__Bifidobacteriaceae_g__Bifidobacterium_s__longum | 3 |
| k__Bacteria_p__Firmicutes_c__Clostridia_o__Clostridiales_f__Clostridiaceae | 3 |
| k__Bacteria_p__Firmicutes_c__Clostridia_o__Clostridiales_f__Clostridiaceae_g__SMB53 | 3 |
| k__Bacteria_p__Firmicutes_c__Clostridia_o__Clostridiales_f__Clostridiaceae_g__SMB53_s__ | 3 |
| k__Bacteria_p__Firmicutes_c__Clostridia_o__Clostridiales_f__Veillonellaceae | 3 |
| k__Bacteria_p__Proteobacteria_c__Alphaproteobacteria_o__RF32 | 3 |
| k__Bacteria_p__Proteobacteria_c__Alphaproteobacteria_o__RF32_f__ | 3 |
| k__Bacteria_p__Proteobacteria_c__Alphaproteobacteria_o__RF32_f___g__ | 3 |

| | |
|---|---|
| k__Bacteria_p__Proteobacteria_c__Alphaproteobacteria_o__RF32_f___g___s__ | 3 |
| k__Bacteria_p__Proteobacteria_c__Deltaproteobacteria | 3 |
| k__Bacteria_p__Proteobacteria_c__Deltaproteobacteria_o__Desulfovibrionales | 3 |
| k__Bacteria_p__Proteobacteria_c__Deltaproteobacteria_o__Desulfovibrionales_f__Desulfovibrionaceae | 3 |
| k__Bacteria_p__Proteobacteria_c__Deltaproteobacteria_o__Desulfovibrionales_f__Desulfovibrionaceae_g__Bilophila | 3 |
| k__Bacteria_p__Proteobacteria_c__Deltaproteobacteria_o__Desulfovibrionales_f__Desulfovibrionaceae_g__Bilophila_s__ | 3 |
| k__Bacteria_p__Verrucomicrobia_c__Verrucomicrobiae | 3 |
| k__Bacteria_p__Verrucomicrobia_c__Verrucomicrobiae_o__Verrucomicrobiales | 3 |
| k__Bacteria_p__Verrucomicrobia_c__Verrucomicrobiae_o__Verrucomicrobiales_f__Verrucomicrobiaceae | 3 |
| k__Bacteria_p__Verrucomicrobia_c__Verrucomicrobiae_o__Verrucomicrobiales_f__Verrucomicrobiaceae_g__Akkermansia | 3 |
| k__Bacteria_p__Verrucomicrobia_c__Verrucomicrobiae_o__Verrucomicrobiales_f__Verrucomicrobiaceae_g__Akkermansia_s__muciniphila | 3 |
| k__Bacteria_p__Actinobacteria_c__Actinobacteria_o__Bifidobacteriales_f__Bifidobacteriaceae_g__Bifidobacterium_s__adolescentis | 2 |
| k__Bacteria_p__Bacteroidetes_c__Bacteroidia_o__Bacteroidales_f__Rikenellaceae | 2 |
| k__Bacteria_p__Firmicutes_c__Clostridia_o__Clostridiales_f__Lachnospiraceae_g__Coprococcus | 2 |
| k__Bacteria_p__Firmicutes_c__Clostridia_o__Clostridiales_f__Lachnospiraceae_g__Coprococcus_s__ | 2 |
| k__Bacteria_p__Firmicutes_c__Clostridia_o__Clostridiales_f__Lachnospiraceae_g__Coprococcus_s__eutactus | 2 |
| k__Bacteria_p__Firmicutes_c__Clostridia_o__Clostridiales_f__Lachnospiraceae_g__Roseburia | 2 |
| k__Bacteria_p__Firmicutes_c__Clostridia_o__Clostridiales_f__Lachnospiraceae_g__Roseburia_s__ | 2 |
| k__Bacteria_p__Firmicutes_c__Clostridia_o__Clostridiales_f__Peptococcaceae | 2 |
| k__Bacteria_p__Firmicutes_c__Clostridia_o__Clostridiales_f__Peptococcaceae_g__rc4_4 | 2 |
| k__Bacteria_p__Firmicutes_c__Clostridia_o__Clostridiales_f__Peptococcaceae_g__rc4_4_s__ | 2 |
| k__Bacteria_p__Firmicutes_c__Clostridia_o__Clostridiales_f__Veillonellaceae_g__Dialister | 2 |
| k__Bacteria_p__Firmicutes_c__Clostridia_o__Clostridiales_f__Veillonellaceae_g__Dialister_s__ | 2 |
| k__Bacteria_p__Actinobacteria_c__Actinobacteria_o__Actinomycetales_f__Actinomycetaceae | 1 |
| k__Bacteria_p__Actinobacteria_c__Actinobacteria_o__Actinomycetales_f__Actinomycetaceae_g__Varibaculum | 1 |
| k__Bacteria_p__Actinobacteria_c__Actinobacteria_o__Actinomycetales_f__Actinomycetaceae_g__Varibaculum_s__ | 1 |
| k__Bacteria_p__Bacteroidetes_c__Bacteroidia_o__Bacteroidales_f__Bacteroidaceae_g__Bacteroides_s__eggerthii | 1 |
| k__Bacteria_p__Bacteroidetes_c__Bacteroidia_o__Bacteroidales_f__Bacteroidaceae_g__Bacteroides_s__uniformis | 1 |
| k__Bacteria_p__Bacteroidetes_c__Bacteroidia_o__Bacteroidales_f___Odoribacteraceae_ | 1 |
| k__Bacteria_p__Bacteroidetes_c__Bacteroidia_o__Bacteroidales_f__Rikenellaceae_g__ | 1 |
| k__Bacteria_p__Bacteroidetes_c__Bacteroidia_o__Bacteroidales_f__Rikenellaceae_g___s__ | 1 |
| k__Bacteria_p__Firmicutes_c__Bacilli_o__Bacillales_f__Staphylococcaceae | 1 |
| k__Bacteria_p__Firmicutes_c__Bacilli_o__Bacillales_f__Staphylococcaceae_g__Staphylococcus | 1 |
| k__Bacteria_p__Firmicutes_c__Bacilli_o__Bacillales_f__Staphylococcaceae_g__Staphylococcus_s__ | 1 |

| | |
|---|---|
| k__Bacteria_p__Firmicutes_c__Bacilli_o__Lactobacillales_f__Lactobacillaceae | 1 |
| k__Bacteria_p__Firmicutes_c__Bacilli_o__Lactobacillales_f__Lactobacillaceae_g__Lactobacillus | 1 |
| k__Bacteria_p__Firmicutes_c__Bacilli_o__Lactobacillales_f__Lactobacillaceae_g__Lactobacillus_s__ | 1 |
| k__Bacteria_p__Firmicutes_c__Bacilli_o__Turicibacterales | 1 |
| k__Bacteria_p__Firmicutes_c__Bacilli_o__Turicibacterales_f__Turicibacteraceae | 1 |
| k__Bacteria_p__Firmicutes_c__Bacilli_o__Turicibacterales_f__Turicibacteraceae_g__Turicibacter | 1 |
| k__Bacteria_p__Firmicutes_c__Bacilli_o__Turicibacterales_f__Turicibacteraceae_g__Turicibacter_s__ | 1 |
| k__Bacteria_p__Firmicutes_c__Clostridia_o__Clostridiales_f__Lachnospiraceae_g__Blautia_s__obeum | 1 |
| k__Bacteria_p__Firmicutes_c__Clostridia_o__Clostridiales_f__Lachnospiraceae_g__Blautia_s__producta | 1 |
| k__Bacteria_p__Firmicutes_c__Clostridia_o__Clostridiales_f__Lachnospiraceae_g__Dorea | 1 |
| k__Bacteria_p__Firmicutes_c__Clostridia_o__Clostridiales_f__Lachnospiraceae_g__Dorea_s__ | 1 |
| k__Bacteria_p__Firmicutes_c__Clostridia_o__Clostridiales_f__Lachnospiraceae_g__Dorea_s__formicigenerans | 1 |
| k__Bacteria_p__Firmicutes_c__Clostridia_o__Clostridiales_f__Lachnospiraceae_g___Ruminococcus__s__gnavus | 1 |
| k__Bacteria_p__Firmicutes_c__Clostridia_o__Clostridiales_f___Mogibacteriaceae_ | 1 |
| k__Bacteria_p__Firmicutes_c__Clostridia_o__Clostridiales_f___Mogibacteriaceae__g__ | 1 |
| k__Bacteria_p__Firmicutes_c__Clostridia_o__Clostridiales_f___Mogibacteriaceae__g___s__ | 1 |
| k__Bacteria_p__Firmicutes_c__Clostridia_o__Clostridiales_f__Peptostreptococcaceae | 1 |
| k__Bacteria_p__Firmicutes_c__Clostridia_o__Clostridiales_f__Peptostreptococcaceae_g__ | 1 |
| k__Bacteria_p__Firmicutes_c__Clostridia_o__Clostridiales_f__Peptostreptococcaceae_g___s__ | 1 |
| k__Bacteria_p__Firmicutes_c__Clostridia_o__Clostridiales_f__Ruminococcaceae | 1 |
| k__Bacteria_p__Firmicutes_c__Clostridia_o__Clostridiales_f__Ruminococcaceae_g__Oscillospira | 1 |
| k__Bacteria_p__Firmicutes_c__Clostridia_o__Clostridiales_f__Ruminococcaceae_g__Oscillospira_s__ | 1 |
| k__Bacteria_p__Firmicutes_c__Clostridia_o__Clostridiales_f___Tissierellaceae__g__Peptoniphilus | 1 |
| k__Bacteria_p__Firmicutes_c__Clostridia_o__Clostridiales_f___Tissierellaceae__g__Peptoniphilus_s__ | 1 |
| k__Bacteria_p__Firmicutes_c__Clostridia_o__Clostridiales_f___Tissierellaceae__g__WAL_1855D | 1 |
| k__Bacteria_p__Firmicutes_c__Clostridia_o__Clostridiales_f___Tissierellaceae__g__WAL_1855D_s__ | 1 |
| k__Bacteria_p__Firmicutes_c__Clostridia_o__Clostridiales_f__Veillonellaceae_g__Acidaminococcus | 1 |
| k__Bacteria_p__Firmicutes_c__Clostridia_o__Clostridiales_f__Veillonellaceae_g__Acidaminococcus_s__ | 1 |
| k__Bacteria_p__Firmicutes_c__Clostridia_o__Clostridiales_f__Veillonellaceae_g__Phascolarctobacterium | 1 |
| k__Bacteria_p__Firmicutes_c__Clostridia_o__Clostridiales_f__Veillonellaceae_g__Phascolarctobacterium_s__ | 1 |
| k__Bacteria_p__Firmicutes_c__Clostridia_o__Clostridiales_f__Veillonellaceae_g__Veillonella | 1 |
| k__Bacteria_p__Firmicutes_c__Clostridia_o__Clostridiales_f__Veillonellaceae_g__Veillonella_s__dispar | 1 |
| k__Bacteria_p__Firmicutes_c__Erysipelotrichi | 1 |
| k__Bacteria_p__Firmicutes_c__Erysipelotrichi_o__Erysipelotrichales | 1 |

| | |
|---|---|
| k__Bacteria_p__Firmicutes_c__Erysipelotrichi_o__Erysipelotrichales_f__Erysipelotrichaceae | 1 |
| k__Bacteria_p__Firmicutes_c__Erysipelotrichi_o__Erysipelotrichales_f__Erysipelotrichaceae_g__ | 1 |
| k__Bacteria_p__Firmicutes_c__Erysipelotrichi_o__Erysipelotrichales_f__Erysipelotrichaceae_g___s__ | 1 |
| k__Bacteria_p__Proteobacteria_c__Alphaproteobacteria | 1 |
| k__Bacteria_p__Proteobacteria_c__Deltaproteobacteria_o__Desulfovibrionales_f__Desulfovibrionaceae_g__Desulfovibrio | 1 |
| k__Bacteria_p__Proteobacteria_c__Gammaproteobacteria_o__Enterobacteriales_f__Enterobacteriaceae_g__Klebsiella | 1 |
| k__Bacteria_p__Proteobacteria_c__Gammaproteobacteria_o__Enterobacteriales_f__Enterobacteriaceae_g__Klebsiella_s__ | 1 |
| k__Bacteria_p__Proteobacteria_c__Gammaproteobacteria_o__Oceanospirillales | 1 |
| k__Bacteria_p__Proteobacteria_c__Gammaproteobacteria_o__Oceanospirillales_f__Halomonadaceae | 1 |

**Supplementary Table 4:** KEGG pathway analysis according to *HLA-B27* status across each sampling site. Associations are color-coded in the following manner: occurred in six sites (red), occurred in five sites (green), occurred in four sites (yellow), occurred in three sites (blue), and occurred in two sites (orange).

| ID | KEGG Pathway | HLA Type | Coefficient | P-value | Q-value | Site |
|---|---|---|---|---|---|---|
| ko00040 | Pentose and glucuronate interconversions | B27 Pos | -0.0054 | 0.0286 | 0.2169 | Ileum |
| ko00052 | Galactose metabolism | B27 Pos | -0.0067 | 0.0059 | 0.2158 | Ileum |
| ko00061 | Fatty acid biosynthesis | B27 Pos | -0.0018 | 0.0223 | 0.1705 | RC |
| ko00071 | Fatty acid metabolism | B27 Pos | 0.0065 | 0.0302 | 0.2169 | Ileum |
| ko00121 | Secondary bile acid biosynthesis | B27 Pos | -0.0116 | 0.0044 | 0.2158 | Ileum |
| ko00121 | Secondary bile acid biosynthesis | B27 Pos | 0.0079 | 0.0439 | 0.6657 | Stool |
| ko00130 | Ubiquinone and other terpenoid-quinone biosynthesis | B27 Pos | 0.0097 | 0.0147 | 0.2158 | Ileum |
| ko00130 | Ubiquinone and other terpenoid-quinone biosynthesis | B27 Pos | 0.0087 | 0.0222 | 0.1705 | RC |
| ko00140 | Steroid hormone biosynthesis | B27 Pos | -0.0050 | 0.0225 | 0.2158 | Ileum |
| ko00140 | Steroid hormone biosynthesis | B27 Pos | -0.0039 | 0.0325 | 0.3860 | LC |
| ko00190 | Oxidative phosphorylation | B27 Pos | -0.0011 | 0.0480 | 0.2398 | RC |
| ko00230 | Purine metabolism | B27 Pos | -0.0012 | 0.0126 | 0.1623 | RC |
| ko00230 | Purine metabolism | B27 Pos | -0.0008 | 0.0337 | 0.6657 | Stool |
| ko00240 | Pyrimidine metabolism | B27 Pos | -0.0021 | 0.0111 | 0.3100 | LC |
| ko00240 | Pyrimidine metabolism | B27 Pos | -0.0019 | 0.0254 | 0.1736 | RC |
| ko00250 | Alanine | B27 Pos | -0.0028 | 0.0201 | 0.3100 | LC |
| ko00250 | Alanine | B27 Pos | -0.0028 | 0.0242 | 0.1736 | RC |
| ko00250 | Alanine | B27 Pos | -0.0031 | 0.0259 | 0.2169 | Ileum |
| ko00260 | Glycine | B27 Pos | 0.0012 | 0.0355 | 0.2177 | Ileum |
| ko00281 | Geraniol degradation | B27 Pos | -0.0143 | 0.0416 | 0.6344 | Cecum |
| ko00290 | Valine | B27 Pos | -0.0036 | 0.0203 | 0.1705 | RC |
| ko00300 | Lysine biosynthesis | B27 Pos | -0.0031 | 0.0232 | 0.3100 | LC |
| ko00300 | Lysine biosynthesis | B27 Pos | -0.0030 | 0.0470 | 0.2398 | RC |
| ko00310 | Lysine degradation | B27 Pos | 0.0054 | 0.0332 | 0.2177 | Ileum |
| ko00310 | Lysine degradation | B27 Pos | 0.0017 | 0.0378 | 0.9890 | Rectum |
| ko00310 | Lysine degradation | B27 Pos | -0.0008 | 0.0464 | 0.6657 | Stool |
| ko00312 | beta-Lactam resistance | B27 Pos | -0.0034 | 0.0411 | 0.3915 | LC |
| ko00330 | Arginine and proline metabolism | B27 Pos | -0.0011 | 0.0470 | 0.3915 | LC |
| ko00340 | Histidine metabolism | B27 Pos | -0.0022 | 0.0191 | 0.3100 | LC |

| | | | | | | |
|---|---|---|---|---|---|---|
| ko00340 | Histidine metabolism | B27 Pos | -0.0030 | 0.0393 | 0.2177 | Ileum |
| ko00380 | Tryptophan metabolism | B27 Pos | 0.0059 | 0.0470 | 0.2177 | Ileum |
| ko00440 | Phosphonate and phosphinate metabolism | B27 Pos | 0.0031 | 0.0204 | 0.5030 | Stool |
| ko00460 | Cyanoamino acid metabolism | B27 Pos | -0.0297 | 0.0195 | 0.5394 | Cecum |
| ko00471 | D-Glutamine and D-glutamate metabolism | B27 Pos | -0.0033 | 0.0291 | 0.1837 | RC |
| ko00473 | D-Alanine metabolism | B27 Pos | 0.0042 | 0.0352 | 0.6344 | Cecum |
| ko00473 | D-Alanine metabolism | B27 Pos | 0.0052 | 0.0372 | 0.2177 | Ileum |
| ko00480 | Glutathione metabolism | B27 Pos | 0.0078 | 0.0148 | 0.2158 | Ileum |
| ko00500 | Starch and sucrose metabolism | B27 Pos | -0.0056 | 0.0106 | 0.2158 | Ileum |
| ko00511 | Other glycan degradation | B27 Pos | -0.0194 | 0.0451 | 0.2177 | Ileum |
| ko00521 | Streptomycin biosynthesis | B27 Pos | -0.0074 | 0.0227 | 0.2158 | Ileum |
| ko00523 | Polyketide sugar unit biosynthesis | B27 Pos | -0.0059 | 0.0134 | 0.2158 | Ileum |
| ko00550 | Peptidoglycan biosynthesis | B27 Pos | -0.0033 | 0.0099 | 0.1426 | RC |
| ko00562 | Inositol phosphate metabolism | B27 Pos | 0.0025 | 0.0009 | 0.0860 | RC |
| ko00562 | Inositol phosphate metabolism | B27 Pos | 0.0016 | 0.0181 | 0.3100 | LC |
| ko00600 | Sphingolipid metabolism | B27 Pos | -0.0164 | 0.0370 | 0.6344 | Cecum |
| ko00620 | Pyruvate metabolism | B27 Pos | -0.0011 | 0.0373 | 0.2205 | RC |
| ko00625 | Chloroalkane and chloroalkene degradation | B27 Pos | -0.0235 | 0.0186 | 0.5394 | Cecum |
| ko00626 | Naphthalene degradation | B27 Pos | 0.0208 | 0.0179 | 0.5394 | Cecum |
| ko00650 | Butanoate metabolism | B27 Pos | 0.0038 | 0.0470 | 0.2177 | Ileum |
| ko00670 | One carbon pool by folate | B27 Pos | -0.0021 | 0.0485 | 0.3915 | LC |
| ko00710 | Carbon fixation in photosynthetic organisms | B27 Pos | -0.0015 | 0.0062 | 0.4744 | Stool |
| ko00710 | Carbon fixation in photosynthetic organisms | B27 Pos | -0.0026 | 0.0424 | 0.2395 | RC |
| ko00720 | Carbon fixation pathways in prokaryotes | B27 Pos | -0.0024 | 0.0177 | 0.5030 | Stool |
| ko00730 | Thiamine metabolism | B27 Pos | -0.0043 | 0.0210 | 0.1705 | RC |
| ko00760 | Nicotinate and nicotinamide metabolism | B27 Pos | -0.0015 | 0.0130 | 0.5030 | Stool |
| ko00770 | Pantothenate and CoA biosynthesis | B27 Pos | -0.0027 | 0.0150 | 0.1623 | RC |
| ko00830 | Retinol metabolism | B27 Pos | 0.0038 | 0.0059 | 0.1268 | RC |
| ko00830 | Retinol metabolism | B27 Pos | 0.0028 | 0.0446 | 0.6344 | Cecum |
| ko00860 | Porphyrin and chlorophyll metabolism | B27 Pos | -0.0032 | 0.0453 | 0.2398 | RC |
| ko00900 | Terpenoid backbone biosynthesis | B27 Pos | -0.0032 | 0.0097 | 0.1426 | RC |
| ko00908 | Zeatin biosynthesis | B27 Pos | -0.0024 | 0.0191 | 0.3100 | LC |
| ko00908 | Zeatin biosynthesis | B27 Pos | -0.0036 | 0.0411 | 0.2177 | Ileum |

| | | | | | | |
|---|---|---|---|---|---|---|
| ko00910 | Nitrogen metabolism | B27 Pos | 0.0026 | 0.0182 | 0.2158 | Ileum |
| ko00983 | Drug metabolism - other enzymes | B27 Pos | -0.0025 | 0.0139 | 0.1623 | RC |
| ko00983 | Drug metabolism - other enzymes | B27 Pos | -0.0029 | 0.0218 | 0.2158 | Ileum |
| ko01040 | Biosynthesis of unsaturated fatty acids | B27 Pos | 0.0088 | 0.0138 | 0.2158 | Ileum |
| ko01040 | Biosynthesis of unsaturated fatty acids | B27 Pos | 0.0025 | 0.0296 | 0.9890 | Rectum |
| ko01053 | Biosynthesis of siderophore group nonribosomal peptides | B27 Pos | 0.0068 | 0.0076 | 0.1411 | RC |
| ko01053 | Biosynthesis of siderophore group nonribosomal peptides | B27 Pos | 0.0073 | 0.0194 | 0.2158 | Ileum |
| ko02010 | ABC transporters | B27 Pos | 0.0040 | 0.0466 | 0.2177 | Ileum |
| ko02020 | Two-component system | B27 Pos | 0.0058 | 0.0178 | 0.2158 | Ileum |
| ko02030 | Bacterial chemotaxis | B27 Pos | 0.0079 | 0.0378 | 0.9890 | Rectum |
| ko02040 | Flagellar assembly | B27 Pos | 0.0098 | 0.0120 | 0.5394 | Cecum |
| ko02040 | Flagellar assembly | B27 Pos | 0.0099 | 0.0161 | 0.3100 | LC |
| ko02040 | Flagellar assembly | B27 Pos | 0.0101 | 0.0199 | 0.1705 | RC |
| ko02040 | Flagellar assembly | B27 Pos | 0.0093 | 0.0219 | 0.9890 | Rectum |
| ko02040 | Flagellar assembly | B27 Pos | 0.0107 | 0.0310 | 0.2169 | Ileum |
| ko03020 | RNA polymerase | B27 Pos | -0.0084 | 0.0007 | 0.0930 | Ileum |
| ko03020 | RNA polymerase | B27 Pos | -0.0021 | 0.0237 | 0.3100 | LC |
| ko03030 | DNA replication | B27 Pos | -0.0025 | 0.0009 | 0.1126 | LC |
| ko03030 | DNA replication | B27 Pos | -0.0027 | 0.0013 | 0.0860 | RC |
| ko03060 | Protein export | B27 Pos | -0.0017 | 0.0297 | 0.1837 | RC |
| ko03060 | Protein export | B27 Pos | -0.0013 | 0.0429 | 0.3915 | LC |
| ko03410 | Base excision repair | B27 Pos | -0.0016 | 0.0057 | 0.1268 | RC |
| ko03410 | Base excision repair | B27 Pos | -0.0017 | 0.0077 | 0.4744 | Stool |
| ko03440 | Homologous recombination | B27 Pos | -0.0027 | 0.0051 | 0.1268 | RC |
| ko03440 | Homologous recombination | B27 Pos | -0.0022 | 0.0136 | 0.3100 | LC |
| ko04112 | Cell cycle - Caulobacter | B27 Pos | -0.0035 | 0.0042 | 0.1268 | RC |
| ko04122 | Sulfur relay system | B27 Pos | 0.0045 | 0.0265 | 0.2169 | Ileum |
| ko04621 | NOD-like receptor signaling pathway | B27 Pos | -0.0011 | 0.0211 | 0.5394 | Cecum |
| ko05100 | Bacterial invasion of epithelial cells | B27 Pos | 0.0028 | 0.0480 | 0.2177 | Ileum |
| ko05120 | Epithelial cell signaling in Helicobacter pylori infection | B27 Pos | -0.0013 | 0.0354 | 0.3860 | LC |
| ko05120 | Epithelial cell signaling in Helicobacter pylori infection | B27 Pos | -0.0012 | 0.0499 | 0.2177 | Ileum |

| ID | KEGG Pathway | Number of tissue sites |
|---|---|---|
| ko02040 | Flagellar assembly | 5 |
| ko00250 | Alanine | 3 |
| ko00310 | Lysine degradation | 3 |
| ko00121 | Secondary bile acid biosynthesis | 2 |
| ko00130 | Ubiquinone and other terpenoid-quinone biosynthesis | 2 |
| ko00140 | Steroid hormone biosynthesis | 2 |
| ko00230 | Purine metabolism | 2 |
| ko00240 | Pyrimidine metabolism | 2 |
| ko00300 | Lysine biosynthesis | 2 |
| ko00340 | Histidine metabolism | 2 |
| ko00473 | D-Alanine metabolism | 2 |
| ko00562 | Inositol phosphate metabolism | 2 |
| ko00710 | Carbon fixation in photosynthetic organisms | 2 |
| ko00830 | Retinol metabolism | 2 |
| ko00908 | Zeatin biosynthesis | 2 |
| ko00983 | Drug metabolism - other enzymes | 2 |
| ko01040 | Biosynthesis of unsaturated fatty acids | 2 |
| ko01053 | Biosynthesis of siderophore group nonribosomal peptides | 2 |
| ko03020 | RNA polymerase | 2 |
| ko03030 | DNA replication | 2 |
| ko03060 | Protein export | 2 |
| ko03410 | Base excision repair | 2 |
| ko03440 | Homologous recombination | 2 |
| ko05120 | Epithelial cell signaling in Helicobacter pylori infection | 2 |
| ko00040 | Pentose and glucuronate interconversions | 1 |
| ko00052 | Galactose metabolism | 1 |
| ko00061 | Fatty acid biosynthesis | 1 |
| ko00071 | Fatty acid metabolism | 1 |
| ko00190 | Oxidative phosphorylation | 1 |
| ko00260 | Glycine | 1 |
| ko00281 | Geraniol degradation | 1 |
| ko00290 | Valine | 1 |
| ko00312 | beta-Lactam resistance | 1 |

| | | |
|---|---|---|
| ko00330 | Arginine and proline metabolism | 1 |
| ko00380 | Tryptophan metabolism | 1 |
| ko00440 | Phosphonate and phosphinate metabolism | 1 |
| ko00460 | Cyanoamino acid metabolism | 1 |
| ko00471 | D-Glutamine and D-glutamate metabolism | 1 |
| ko00480 | Glutathione metabolism | 1 |
| ko00500 | Starch and sucrose metabolism | 1 |
| ko00511 | Other glycan degradation | 1 |
| ko00521 | Streptomycin biosynthesis | 1 |
| ko00523 | Polyketide sugar unit biosynthesis | 1 |
| ko00550 | Peptidoglycan biosynthesis | 1 |
| ko00600 | Sphingolipid metabolism | 1 |
| ko00620 | Pyruvate metabolism | 1 |
| ko00625 | Chloroalkane and chloroalkene degradation | 1 |
| ko00626 | Naphthalene degradation | 1 |
| ko00650 | Butanoate metabolism | 1 |
| ko00670 | One carbon pool by folate | 1 |
| ko00720 | Carbon fixation pathways in prokaryotes | 1 |
| ko00730 | Thiamine metabolism | 1 |
| ko00760 | Nicotinate and nicotinamide metabolism | 1 |
| ko00770 | Pantothenate and CoA biosynthesis | 1 |
| ko00860 | Porphyrin and chlorophyll metabolism | 1 |
| ko00900 | Terpenoid backbone biosynthesis | 1 |
| ko00910 | Nitrogen metabolism | 1 |
| ko02010 | ABC transporters | 1 |
| ko02020 | Two-component system | 1 |
| ko02030 | Bacterial chemotaxis | 1 |
| ko04112 | Cell cycle - Caulobacter | 1 |
| ko04122 | Sulfur relay system | 1 |
| ko04621 | NOD-like receptor signaling pathway | 1 |
| ko05100 | Bacterial invasion of epithelial cells | 1 |

**Supplementary Table 5:** KEGG pathway analysis according to *HLA-DRB1* status across each sampling site. Enrichment or depletion is measured relative to the neutral risk genotype. Associations are color-coded in the following manner: occurred in six sites (red), occurred in five sites (green), occurred in four sites (yellow), occurred in three sites (blue), and occurred in two sites (orange).

| ID | KEGG Pathway | DRB1 type | Coefficient | P-value | Q-value | Site |
|---|---|---|---|---|---|---|
| ko00010 | Glycolysis / Gluconeogenesis | Risk | 0.0008 | 0.0090 | 0.2265 | LC |
| ko00020 | Citrate cycle (TCA cycle) | Risk | -0.0032 | 0.0008 | 0.3392 | Ileum |
| ko00020 | Citrate cycle (TCA cycle) | Risk | -0.0031 | 0.0017 | 0.0886 | LC |
| ko00020 | Citrate cycle (TCA cycle) | Risk | -0.0024 | 0.0050 | 0.5186 | Rectum |
| ko00020 | Citrate cycle (TCA cycle) | Risk | -0.0024 | 0.0078 | 0.4736 | RC |
| ko00020 | Citrate cycle (TCA cycle) | Risk | -0.0021 | 0.0237 | 0.6654 | Cecum |
| ko00030 | Pentose phosphate pathway | Risk | 0.0018 | 0.0110 | 0.2313 | LC |
| ko00030 | Pentose phosphate pathway | Risk | 0.0018 | 0.0186 | 0.9880 | Ileum |
| ko00072 | Synthesis and degradation of ketone bodies | Risk | 0.0070 | 0.0101 | 0.4736 | RC |
| ko00072 | Synthesis and degradation of ketone bodies | Protective | 0.0047 | 0.0380 | 1.0000 | Stool |
| ko00120 | Primary bile acid biosynthesis | Risk | 0.0071 | 0.0171 | 0.7530 | Rectum |
| ko00120 | Primary bile acid biosynthesis | Protective | 0.0113 | 0.0450 | 0.7513 | Cecum |
| ko00130 | Ubiquinone and other terpenoid-quinone biosynthesis | Risk | -0.0043 | 0.0011 | 0.2242 | Rectum |
| ko00140 | Steroid hormone biosynthesis | Risk | -0.0027 | 0.0205 | 0.3350 | LC |
| ko00140 | Steroid hormone biosynthesis | Risk | -0.0023 | 0.0378 | 0.7513 | Cecum |
| ko00140 | Steroid hormone biosynthesis | Risk | -0.0022 | 0.0401 | 0.8503 | Rectum |
| ko00190 | Oxidative phosphorylation | Risk | -0.0007 | 0.0283 | 0.8452 | RC |
| ko00190 | Oxidative phosphorylation | Risk | -0.0006 | 0.0294 | 1.0000 | Ileum |
| ko00190 | Oxidative phosphorylation | Risk | -0.0008 | 0.0297 | 0.3792 | LC |
| ko00230 | Purine metabolism | Risk | 0.0005 | 0.0264 | 0.3733 | LC |
| ko00230 | Purine metabolism | Risk | 0.0005 | 0.0282 | 1.0000 | Stool |
| ko00260 | Glycine | Risk | 0.0007 | 0.0350 | 1.0000 | Stool |
| ko00270 | Cysteine and methionine metabolism | Risk | 0.0012 | 0.0067 | 0.4736 | RC |
| ko00270 | Cysteine and methionine metabolism | Risk | 0.0011 | 0.0118 | 0.2369 | LC |
| ko00270 | Cysteine and methionine metabolism | Risk | 0.0011 | 0.0214 | 0.7530 | Rectum |
| ko00280 | Valine | Risk | -0.0016 | 0.0486 | 0.8503 | Rectum |
| ko00281 | Geraniol degradation | Risk | -0.0108 | 0.0377 | 1.0000 | Ileum |
| ko00300 | Lysine biosynthesis | Risk | 0.0025 | 0.0019 | 0.0886 | LC |

| | | | | | | |
|---|---|---|---|---|---|---|
| ko00310 | Lysine degradation | Risk | -0.0011 | 0.0287 | 0.7973 | Rectum |
| ko00311 | Penicillin and cephalosporin biosynthesis | Risk | -0.0037 | 0.0015 | 0.0886 | LC |
| ko00311 | Penicillin and cephalosporin biosynthesis | Risk | -0.0033 | 0.0028 | 0.3549 | Cecum |
| ko00311 | Penicillin and cephalosporin biosynthesis | Risk | -0.0030 | 0.0085 | 0.4736 | RC |
| ko00311 | Penicillin and cephalosporin biosynthesis | Risk | -0.0035 | 0.0089 | 1.0000 | Stool |
| ko00311 | Penicillin and cephalosporin biosynthesis | Risk | -0.0028 | 0.0184 | 0.7530 | Rectum |
| ko00362 | Benzoate degradation | Risk | 0.0012 | 0.0171 | 1.0000 | Stool |
| ko00362 | Benzoate degradation | Risk | 0.0011 | 0.0391 | 0.4271 | LC |
| ko00380 | Tryptophan metabolism | Protective | -0.0090 | 0.0452 | 0.7513 | Cecum |
| ko00400 | Phenylalanine | Risk | 0.0016 | 0.0201 | 0.6654 | Cecum |
| ko00400 | Phenylalanine | Protective | 0.0025 | 0.0300 | 0.7047 | Cecum |
| ko00400 | Phenylalanine | Risk | 0.0015 | 0.0309 | 0.7973 | Rectum |
| ko00400 | Phenylalanine | Risk | 0.0016 | 0.0363 | 0.4167 | LC |
| ko00430 | Taurine and hypotaurine metabolism | Risk | -0.0010 | 0.0130 | 0.2522 | LC |
| ko00430 | Taurine and hypotaurine metabolism | Risk | -0.0010 | 0.0215 | 0.6654 | Cecum |
| ko00440 | Phosphonate and phosphinate metabolism | Risk | -0.0020 | 0.0143 | 0.2579 | LC |
| ko00450 | Selenocompound metabolism | Protective | 0.0017 | 0.0305 | 0.7047 | Cecum |
| ko00480 | Glutathione metabolism | Risk | -0.0035 | 0.0099 | 0.2265 | LC |
| ko00500 | Starch and sucrose metabolism | Risk | 0.0016 | 0.0426 | 0.4331 | LC |
| ko00510 | N-Glycan biosynthesis | Risk | -0.0014 | 0.0084 | 0.2265 | LC |
| ko00510 | N-Glycan biosynthesis | Risk | -0.0012 | 0.0205 | 0.7530 | Rectum |
| ko00510 | N-Glycan biosynthesis | Risk | -0.0011 | 0.0375 | 0.9225 | RC |
| ko00511 | Other glycan degradation | Risk | -0.0085 | 0.0447 | 0.8503 | Rectum |
| ko00531 | Glycosaminoglycan degradation | Risk | -0.0090 | 0.0082 | 0.4736 | RC |
| ko00531 | Glycosaminoglycan degradation | Risk | -0.0090 | 0.0108 | 0.2313 | LC |
| ko00531 | Glycosaminoglycan degradation | Risk | -0.0073 | 0.0232 | 0.7530 | Rectum |
| ko00531 | Glycosaminoglycan degradation | Risk | -0.0074 | 0.0249 | 0.6654 | Cecum |
| ko00531 | Glycosaminoglycan degradation | Risk | -0.0073 | 0.0290 | 1.0000 | Stool |
| ko00540 | Lipopolysaccharide biosynthesis | Risk | -0.0081 | 0.0012 | 0.4651 | RC |
| ko00540 | Lipopolysaccharide biosynthesis | Risk | -0.0075 | 0.0013 | 0.2242 | Rectum |
| ko00540 | Lipopolysaccharide biosynthesis | Risk | -0.0079 | 0.0020 | 0.0887 | LC |
| ko00540 | Lipopolysaccharide biosynthesis | Risk | -0.0077 | 0.0030 | 0.3901 | Ileum |
| ko00540 | Lipopolysaccharide biosynthesis | Risk | -0.0062 | 0.0178 | 0.6654 | Cecum |

| | | | | | | |
|---|---|---|---|---|---|---|
| ko00550 | Peptidoglycan biosynthesis | Risk | 0.0017 | 0.0312 | 0.3895 | LC |
| ko00561 | Glycerolipid metabolism | Risk | 0.0022 | 0.0003 | 0.0438 | LC |
| ko00561 | Glycerolipid metabolism | Risk | 0.0024 | 0.0013 | 0.3392 | Ileum |
| ko00561 | Glycerolipid metabolism | Risk | 0.0016 | 0.0092 | 0.4736 | RC |
| ko00561 | Glycerolipid metabolism | Risk | 0.0016 | 0.0184 | 0.6654 | Cecum |
| ko00561 | Glycerolipid metabolism | Risk | 0.0016 | 0.0233 | 0.7530 | Rectum |
| ko00562 | Inositol phosphate metabolism | Risk | -0.0012 | 0.0070 | 0.6066 | Cecum |
| ko00562 | Inositol phosphate metabolism | Risk | -0.0012 | 0.0079 | 0.6680 | Rectum |
| ko00562 | Inositol phosphate metabolism | Risk | -0.0013 | 0.0124 | 0.5240 | RC |
| ko00562 | Inositol phosphate metabolism | Risk | -0.0010 | 0.0237 | 0.3646 | LC |
| ko00562 | Inositol phosphate metabolism | Protective | -0.0015 | 0.0494 | 0.7513 | Cecum |
| ko00564 | Glycerophospholipid metabolism | Risk | 0.0021 | 0.0004 | 0.0438 | LC |
| ko00564 | Glycerophospholipid metabolism | Risk | 0.0015 | 0.0072 | 0.6066 | Cecum |
| ko00564 | Glycerophospholipid metabolism | Risk | 0.0013 | 0.0232 | 0.8139 | RC |
| ko00564 | Glycerophospholipid metabolism | Risk | 0.0016 | 0.0370 | 1.0000 | Ileum |
| ko00564 | Glycerophospholipid metabolism | Risk | 0.0012 | 0.0483 | 0.8503 | Rectum |
| ko00565 | Ether lipid metabolism | Risk | 0.0021 | 0.0210 | 1.0000 | Ileum |
| ko00620 | Pyruvate metabolism | Risk | 0.0007 | 0.0098 | 0.2265 | LC |
| ko00620 | Pyruvate metabolism | Risk | 0.0008 | 0.0185 | 0.9880 | Ileum |
| ko00627 | Aminobenzoate degradation | Protective | -0.0024 | 0.0238 | 0.6654 | Cecum |
| ko00627 | Aminobenzoate degradation | Risk | -0.0013 | 0.0467 | 0.7513 | Cecum |
| ko00630 | Glyoxylate and dicarboxylate metabolism | Risk | -0.0011 | 0.0091 | 0.6680 | Rectum |
| ko00633 | Nitrotoluene degradation | Risk | 0.0025 | 0.0168 | 0.2929 | LC |
| ko00633 | Nitrotoluene degradation | Risk | 0.0019 | 0.0219 | 0.6654 | Cecum |
| ko00643 | Styrene degradation | Risk | -0.0059 | 0.0290 | 0.3792 | LC |
| ko00670 | One carbon pool by folate | Risk | 0.0015 | 0.0278 | 0.3733 | LC |
| ko00680 | Methane metabolism | Risk | 0.0010 | 0.0467 | 0.4613 | LC |
| ko00720 | Carbon fixation pathways in prokaryotes | Risk | -0.0015 | 0.0127 | 0.8442 | Ileum |
| ko00720 | Carbon fixation pathways in prokaryotes | Risk | -0.0014 | 0.0132 | 0.5240 | RC |
| ko00730 | Thiamine metabolism | Protective | 0.0035 | 0.0100 | 1.0000 | Stool |
| ko00730 | Thiamine metabolism | Risk | 0.0026 | 0.0236 | 0.3646 | LC |
| ko00730 | Thiamine metabolism | Risk | 0.0026 | 0.0301 | 0.7047 | Cecum |
| ko00730 | Thiamine metabolism | Risk | 0.0025 | 0.0333 | 1.0000 | Ileum |

| | | | | | | |
|---|---|---|---|---|---|---|
| ko00730 | Thiamine metabolism | Risk | 0.0022 | 0.0422 | 0.8503 | Rectum |
| ko00730 | Thiamine metabolism | Protective | 0.0040 | 0.0500 | 0.7513 | Cecum |
| ko00750 | Vitamin B6 metabolism | Risk | -0.0016 | 0.0017 | 0.0886 | LC |
| ko00750 | Vitamin B6 metabolism | Risk | -0.0013 | 0.0387 | 0.7513 | Cecum |
| ko00760 | Nicotinate and nicotinamide metabolism | Risk | 0.0010 | 0.0287 | 1.0000 | Stool |
| ko00770 | Pantothenate and CoA biosynthesis | Protective | 0.0027 | 0.0205 | 0.6654 | Cecum |
| ko00780 | Biotin metabolism | Risk | -0.0026 | 0.0036 | 0.4651 | RC |
| ko00780 | Biotin metabolism | Risk | -0.0027 | 0.0067 | 0.2192 | LC |
| ko00780 | Biotin metabolism | Risk | -0.0025 | 0.0189 | 0.7530 | Rectum |
| ko00780 | Biotin metabolism | Risk | -0.0023 | 0.0383 | 1.0000 | Ileum |
| ko00785 | Lipoic acid metabolism | Risk | -0.0060 | 0.0097 | 0.2265 | LC |
| ko00785 | Lipoic acid metabolism | Risk | -0.0053 | 0.0332 | 0.9021 | RC |
| ko00785 | Lipoic acid metabolism | Risk | -0.0050 | 0.0411 | 0.7513 | Cecum |
| ko00785 | Lipoic acid metabolism | Risk | -0.0044 | 0.0494 | 0.8503 | Rectum |
| ko00790 | Folate biosynthesis | Risk | -0.0020 | 0.0042 | 0.3901 | Ileum |
| ko00790 | Folate biosynthesis | Risk | -0.0019 | 0.0263 | 0.3733 | LC |
| ko00790 | Folate biosynthesis | Risk | -0.0016 | 0.0366 | 0.9225 | RC |
| ko00791 | Atrazine degradation | Risk | -0.0027 | 0.0430 | 0.4331 | LC |
| ko00860 | Porphyrin and chlorophyll metabolism | Risk | 0.0029 | 0.0007 | 0.0610 | LC |
| ko00860 | Porphyrin and chlorophyll metabolism | Protective | 0.0044 | 0.0146 | 0.6654 | Cecum |
| ko00860 | Porphyrin and chlorophyll metabolism | Risk | 0.0021 | 0.0399 | 0.9369 | RC |
| ko00910 | Nitrogen metabolism | Risk | -0.0009 | 0.0429 | 0.4331 | LC |
| ko00930 | Caprolactam degradation | Risk | -0.0099 | 0.0002 | 0.0423 | LC |
| ko00930 | Caprolactam degradation | Protective | -0.0095 | 0.0408 | 0.4331 | LC |
| ko00930 | Caprolactam degradation | Risk | -0.0030 | 0.0425 | 1.0000 | Stool |
| ko00970 | Aminoacyl-tRNA biosynthesis | Risk | 0.0022 | 0.0329 | 0.4015 | LC |
| ko00980 | Metabolism of xenobiotics by cytochrome P450 | Protective | 0.0116 | 0.0054 | 0.4121 | Ileum |
| ko01051 | Biosynthesis of ansamycins | Risk | 0.0103 | 0.0010 | 0.0720 | LC |
| ko01051 | Biosynthesis of ansamycins | Risk | 0.0081 | 0.0237 | 0.8139 | RC |
| ko01051 | Biosynthesis of ansamycins | Risk | 0.0062 | 0.0242 | 0.6654 | Cecum |
| ko01051 | Biosynthesis of ansamycins | Risk | 0.0075 | 0.0458 | 1.0000 | Ileum |
| ko01051 | Biosynthesis of ansamycins | Risk | 0.0056 | 0.0466 | 0.8503 | Rectum |
| ko01053 | Biosynthesis of siderophore group nonribosomal peptides | Risk | -0.0027 | 0.0276 | 0.3733 | LC |

| | | | | | | |
|---|---|---|---|---|---|---|
| ko02010 | ABC transporters | Risk | 0.0031 | 0.0022 | 0.3549 | Cecum |
| ko02010 | ABC transporters | Risk | 0.0029 | 0.0027 | 0.4651 | RC |
| ko02010 | ABC transporters | Risk | 0.0032 | 0.0032 | 0.1207 | LC |
| ko02010 | ABC transporters | Risk | 0.0035 | 0.0044 | 0.3901 | Ileum |
| ko02010 | ABC transporters | Risk | 0.0023 | 0.0356 | 0.8503 | Rectum |
| ko02030 | Bacterial chemotaxis | Risk | 0.0090 | 0.0002 | 0.0423 | LC |
| ko02030 | Bacterial chemotaxis | Risk | 0.0068 | 0.0009 | 0.2400 | Cecum |
| ko02030 | Bacterial chemotaxis | Risk | 0.0056 | 0.0093 | 0.4736 | RC |
| ko02030 | Bacterial chemotaxis | Risk | 0.0056 | 0.0124 | 0.7530 | Rectum |
| ko02030 | Bacterial chemotaxis | Risk | 0.0056 | 0.0258 | 1.0000 | Ileum |
| ko02040 | Flagellar assembly | Risk | 0.0063 | 0.0261 | 0.3733 | LC |
| ko03013 | RNA transport | Risk | 0.0013 | 0.0000 | 0.0034 | LC |
| ko03013 | RNA transport | Risk | 0.0010 | 0.0008 | 0.2242 | Rectum |
| ko03013 | RNA transport | Risk | 0.0009 | 0.0009 | 0.2400 | Cecum |
| ko03013 | RNA transport | Risk | 0.0009 | 0.0019 | 0.4651 | RC |
| ko03013 | RNA transport | Risk | 0.0010 | 0.0036 | 0.3901 | Ileum |
| ko03018 | RNA degradation | Risk | -0.0007 | 0.0202 | 0.3350 | LC |
| ko03018 | RNA degradation | Risk | -0.0005 | 0.0302 | 0.7973 | Rectum |
| ko03020 | RNA polymerase | Risk | -0.0011 | 0.0361 | 0.4167 | LC |
| ko03420 | Nucleotide excision repair | Risk | 0.0009 | 0.0374 | 0.4167 | LC |
| ko04112 | Cell cycle - Caulobacter | Risk | 0.0016 | 0.0372 | 0.4167 | LC |
| ko04141 | Protein processing in endoplasmic reticulum | Risk | -0.0011 | 0.0096 | 0.2265 | LC |
| ko04141 | Protein processing in endoplasmic reticulum | Risk | -0.0008 | 0.0254 | 0.7701 | Rectum |
| ko04141 | Protein processing in endoplasmic reticulum | Risk | -0.0009 | 0.0254 | 0.8188 | RC |
| ko04141 | Protein processing in endoplasmic reticulum | Risk | -0.0010 | 0.0367 | 1.0000 | Ileum |
| ko04141 | Protein processing in endoplasmic reticulum | Risk | -0.0008 | 0.0488 | 0.7513 | Cecum |
| ko04142 | Lysosome | Risk | -0.0068 | 0.0467 | 0.8503 | Rectum |
| ko04621 | NOD-like receptor signaling pathway | Risk | -0.0005 | 0.0074 | 0.2265 | LC |
| ko04626 | Plant-pathogen interaction | Risk | 0.0013 | 0.0055 | 0.1934 | LC |
| ko04626 | Plant-pathogen interaction | Risk | 0.0012 | 0.0145 | 0.6654 | Cecum |
| ko04626 | Plant-pathogen interaction | Risk | 0.0009 | 0.0377 | 1.0000 | Ileum |
| ko04626 | Plant-pathogen interaction | Risk | 0.0009 | 0.0421 | 0.8503 | Rectum |
| ko04974 | Protein digestion and absorption | Risk | -0.0025 | 0.0140 | 0.2579 | LC |

| | | | | | | |
|---|---|---|---|---|---|---|
| ko04974 | Protein digestion and absorption | Risk | -0.0021 | 0.0171 | 0.7530 | Rectum |
| ko04974 | Protein digestion and absorption | Risk | -0.0022 | 0.0295 | 0.8452 | RC |
| ko04974 | Protein digestion and absorption | Risk | -0.0024 | 0.0402 | 1.0000 | Ileum |
| ko04974 | Protein digestion and absorption | Risk | -0.0020 | 0.0416 | 0.7513 | Cecum |
| ko05120 | Epithelial cell signaling in Helicobacter pylori infection | Protective | 0.0014 | 0.0475 | 0.7513 | Cecum |
| ko05146 | Amoebiasis | Risk | -0.0013 | 0.0019 | 0.2501 | Rectum |
| ko05146 | Amoebiasis | Risk | -0.0012 | 0.0020 | 0.9908 | Stool |
| ko05146 | Amoebiasis | Risk | -0.0014 | 0.0029 | 0.1166 | LC |
| ko05146 | Amoebiasis | Risk | -0.0012 | 0.0100 | 0.6654 | Cecum |
| ko05146 | Amoebiasis | Risk | -0.0012 | 0.0231 | 1.0000 | Ileum |

| ID | KEGG Pathway | Number of tissue sites |
|---|---|---|
| ko00730 | Thiamine metabolism | 6 |
| ko00020 | Citrate cycle (TCA cycle) | 5 |
| ko00311 | Penicillin and cephalosporin biosynthesis | 5 |
| ko00531 | Glycosaminoglycan degradation | 5 |
| ko00540 | Lipopolysaccharide biosynthesis | 5 |
| ko00561 | Glycerolipid metabolism | 5 |
| ko00562 | Inositol phosphate metabolism | 5 |
| ko00564 | Glycerophospholipid metabolism | 5 |
| ko01051 | Biosynthesis of ansamycins | 5 |
| ko02010 | ABC transporters | 5 |
| ko02030 | Bacterial chemotaxis | 5 |
| ko03013 | RNA transport | 5 |
| ko04141 | Protein processing in endoplasmic reticulum | 5 |
| ko04974 | Protein digestion and absorption | 5 |
| ko05146 | Amoebiasis | 5 |
| ko00400 | Phenylalanine | 4 |
| ko00780 | Biotin metabolism | 4 |
| ko00785 | Lipoic acid metabolism | 4 |
| ko04626 | Plant-pathogen interaction | 4 |
| ko00140 | Steroid hormone biosynthesis | 3 |
| ko00190 | Oxidative phosphorylation | 3 |

| | | |
|---|---|---|
| ko00270 | Cysteine and methionine metabolism | 3 |
| ko00510 | N-Glycan biosynthesis | 3 |
| ko00790 | Folate biosynthesis | 3 |
| ko00860 | Porphyrin and chlorophyll metabolism | 3 |
| ko00930 | Caprolactam degradation | 3 |
| ko00030 | Pentose phosphate pathway | 2 |
| ko00072 | Synthesis and degradation of ketone bodies | 2 |
| ko00120 | Primary bile acid biosynthesis | 2 |
| ko00230 | Purine metabolism | 2 |
| ko00362 | Benzoate degradation | 2 |
| ko00430 | Taurine and hypotaurine metabolism | 2 |
| ko00620 | Pyruvate metabolism | 2 |
| ko00627 | Aminobenzoate degradation | 2 |
| ko00633 | Nitrotoluene degradation | 2 |
| ko00720 | Carbon fixation pathways in prokaryotes | 2 |
| ko00750 | Vitamin B6 metabolism | 2 |
| ko03018 | RNA degradation | 2 |
| ko00010 | Glycolysis / Gluconeogenesis | 1 |
| ko00130 | Ubiquinone and other terpenoid-quinone biosynthesis | 1 |
| ko00260 | Glycine | 1 |
| ko00280 | Valine | 1 |
| ko00281 | Geraniol degradation | 1 |
| ko00300 | Lysine biosynthesis | 1 |
| ko00310 | Lysine degradation | 1 |
| ko00380 | Tryptophan metabolism | 1 |
| ko00440 | Phosphonate and phosphinate metabolism | 1 |
| ko00450 | Selenocompound metabolism | 1 |
| ko00480 | Glutathione metabolism | 1 |
| ko00500 | Starch and sucrose metabolism | 1 |
| ko00511 | Other glycan degradation | 1 |
| ko00550 | Peptidoglycan biosynthesis | 1 |
| ko00565 | Ether lipid metabolism | 1 |
| ko00630 | Glyoxylate and dicarboxylate metabolism | 1 |
| ko00643 | Styrene degradation | 1 |

| | | |
|---|---|---|
| ko00670 | One carbon pool by folate | 1 |
| ko00680 | Methane metabolism | 1 |
| ko00760 | Nicotinate and nicotinamide metabolism | 1 |
| ko00770 | Pantothenate and CoA biosynthesis | 1 |
| ko00791 | Atrazine degradation | 1 |
| ko00910 | Nitrogen metabolism | 1 |
| ko00970 | Aminoacyl-tRNA biosynthesis | 1 |
| ko00980 | Metabolism of xenobiotics by cytochrome P450 | 1 |
| ko01053 | Biosynthesis of siderophore group nonribosomal peptides | 1 |
| ko02040 | Flagellar assembly | 1 |
| ko03020 | RNA polymerase | 1 |
| ko03420 | Nucleotide excision repair | 1 |
| ko04112 | Cell cycle - Caulobacter | 1 |
| ko04142 | Lysosome | 1 |
| ko04621 | NOD-like receptor signaling pathway | 1 |
| ko05120 | Epithelial cell signaling in Helicobacter pylori infection | 1 |
